# Genomic stock structure of the marine teleost tarakihi (*Nemadactylus macropterus*) provides evidence of fine-scale adaptation and a temperature-associated cline amid panmixia

**DOI:** 10.1101/2022.02.10.479861

**Authors:** Yvan Papa, Mark A. Morrison, Maren Wellenreuther, Peter A. Ritchie

## Abstract

Tarakihi (*Nemadactylus macropterus*) is an important fishery species with widespread distribution around New Zealand and off the southern coasts of Australia. However, little is known about whether the populations are locally adapted or genetically structured. To address this, we conducted whole-genome resequencing of 175 tarakihi from around New Zealand and Tasmania (Australia) to obtain a dataset of 7.5 million genome-wide and high-quality single nucleotide polymorphisms (SNPs). Variant filtering, *F*_ST_-outlier analysis, and redundancy analysis (RDA) were used to evaluate population structure, adaptive structure, and locus-environment associations. A weak but significant level of neutral genetic differentiation was found between tarakihi from New Zealand and Tasmania (*F*_ST_ = 0.0054–0.0073, *P* ≤ 0.05), supporting the existence of at least two separate reproductive stocks. No clustering was detected among the New Zealand populations (Ф_ST_ < 0.001, *P* = 0.77). Outlier-based, presumably adaptive variation suggests fine-scale adaptive structure between locations around central New Zealand off the east (Wairarapa, Cape Campbell, and Hawke’s Bay) and the west coast (Tasman Bay/Golden Bay and Upper West Coast of South Island). Allele frequencies from 55 loci were associated with at least one of six environmental variables, of which 47 correlated strongly with yearly mean water temperature. Although genes associated with these loci are linked to various functions, the most common functions were integral components of membrane and cilium assembly. Projection of the RDA indicates the existence of a latitudinal temperature cline. Our work provides the first genomic insights supporting panmixia of tarakihi in New Zealand and evidence of a genomic cline that appears to be driven by the temperature gradients, together providing crucial information to inform the stock assessment of this species, and to widen the insights of the ecological drivers of adaptive variation in a marine species.

## 1 Introduction

Effective fisheries management relies on the identification and delineation of stocks to enable optimal and sustainable utilization (Begg et al., 1999; Waples et al., 2008; Cadrin et al., 2014). Ideally, the ultimate goal of fisheries management is to harvest each separate stock at a rate that matches their level of recruitment when taking into account natural mortality (Beddington et al., 2007; Zhou et al., 2019). Discrepancies between the stock management units and the boundaries of biological units can result in overexploitation (Laikre et al., 2005; Reiss et al., 2009; Benestan, 2019), which if left unchecked, could lead to the decline, and ultimately the collapse, of a stock (Orensanz et al., 1998; Ying et al., 2011; Cadrin, 2020). However, biologically accurate information about stock boundaries is still lacking for the vast majority of fisheries species, particularly those residing in the New Zealand Exclusive Economic Zone (Papa et al., 2021b).

Marine environments often contain few physical barriers when compared to freshwater or terrestrial environments. Therefore, the levels of genetic divergence among groups of marine fishes are often expected, and found, to be low (e.g. Koot et al., 2021). This is especially true for marine species with large population sizes and high potential for larval and/or adult dispersal (Ovenden, 2013; Lal et al., 2016; Sandoval-Castillo et al., 2018). Traditional genetic markers (e.g. microsatellites or mitochondrial sequences), that typically represent a very small proportion of the genome, often do not provide the level of resolution required to detect fine-scale genetic structure. However, not detecting any significant genetic differentiation does not necessarily mean there is a level of migration relevant to fisheries management. Populations could be demographically independent but only recently sundered, there could still be some low level of gene flow, or effective population size could be high, with differentiation caused by drift happening slowly (Benestan et al., 2015; Attard et al., 2018). Moreover, when variation is detected with low-resolution markers, it is typically very difficult to determine whether the observed variation is neutral (i.e. due to the accumulation of random mutations through e.g. genetic drift) or adaptive (i.e. due to natural selection which leads to local adaptation) (Carvalho and Hauser, 1994; Allendorf et al., 2010).

In contrast to low-resolution traditional genetic markers, next-generation-sequencing technologies can produce very large genome-wide datasets that have two advantages: (1) a vast increase in the number of available neutral loci, which can meet the level of resolution required when testing for genetic differentiation in marine species, and (2) the ability to detect genomic regions that are currently, or at some time in the recent past, experiencing selection (Bernatchez et al., 2017; Benestan, 2019; Papa et al., 2021b). Previously unknown population structure has been detected in several marine species using genome-wide single nucleotide polymorphism (SNP) datasets. They include the American lobster (Benestan et al., 2015), yellowfin tuna (Pecoraro et al., 2018), silky shark (Kraft et al., 2020), green abalone (Mejía-Ruíz et al., 2020), and California market squid (Cheng et al., 2021). Even when populations display little to no neutral genomic differentiation, adaptive population structure can be detected through outlier-based methods (e.g. albacore (Vaux et al., 2021)) or environment association methods (e.g. American lobster (Benestan et al., 2016), summer flounder (Hoey and Pinsky, 2018), or greenlip abalone (Sandoval-Castillo et al., 2018)).

Tarakihi (*Nemadactylus macropterus* (Forster 1801)) (Figure 1A) is a demersal marine fish species with an expansive distribution, being widely found in the inshore areas of New Zealand (Figure 2). The species occurs from the Three Kings Islands in the north to the Snare Islands in the south and the Chatham Islands in the east, at depths of 10 to 250 m (Annala, 1987; Roberts et al., 2015). It is also distributed along the southern inshore areas of Australia, including Tasmania (Roberts et al., 2015). They are broadcast spawners that form serial breeding aggregations during summer and autumn (Tong and Vooren, 1972). Tarakihi late-stage larvae go through an epipelagic “paperfish” larval phase for approximately 10 months (Annala, 1987; Roberts et al., 2015) (Figure 1B). During this period, their dispersal is mainly driven by oceanic currents, where mixing of individuals from different spawning areas can occur (Bruce, 2001). After c. 10 months, post-larvae morph into juveniles and settle in shallow nursing grounds (Vooren, 1972) (Figure 1C). Adults can live for more than 30 years and have the potential to disperse over large distances, sometimes hundreds of kilometers (Annala, 1987; Hanchet and Field, 2001). Tarakihi are mainly caught by bottom trawling at depths of about 250 m. Commercial catches in New Zealand are around 5,000 tons per year over the past 30 years, with a very recent reduction to 4,400 tons for the fishing year 2019-2020 (Fisheries New Zealand, 2021). While tarakihi are commercially caught in all the unprotected Quota Management Areas of the New Zealand Exclusive Economic Zone, the majority of catches (c. 80%) occur off the east coast of the North and South Island (Langley, 2018). The spawning biomass in some areas (TAR1, TAR2 and TAR3, Figure 2) are thought to be below the fisheries management soft limit (20% of the unexploited, equilibrium biomass) since the early 2000s (Langley, 2018). Consequently, the total allowable commercial catch (TACC) was reduced in 2018 and again in 2019 for these areas, which is the first reduction in tarakihi TACC since the 1980s (Fisheries New Zealand, 2021). The observed declines highlight the need for evidence-based management strategies that incorporate best knowledge about the biological stock structure of this species. However, both stock structure and the levels of connectivity among fished areas are poorly known for this species.

**Figure 1.**
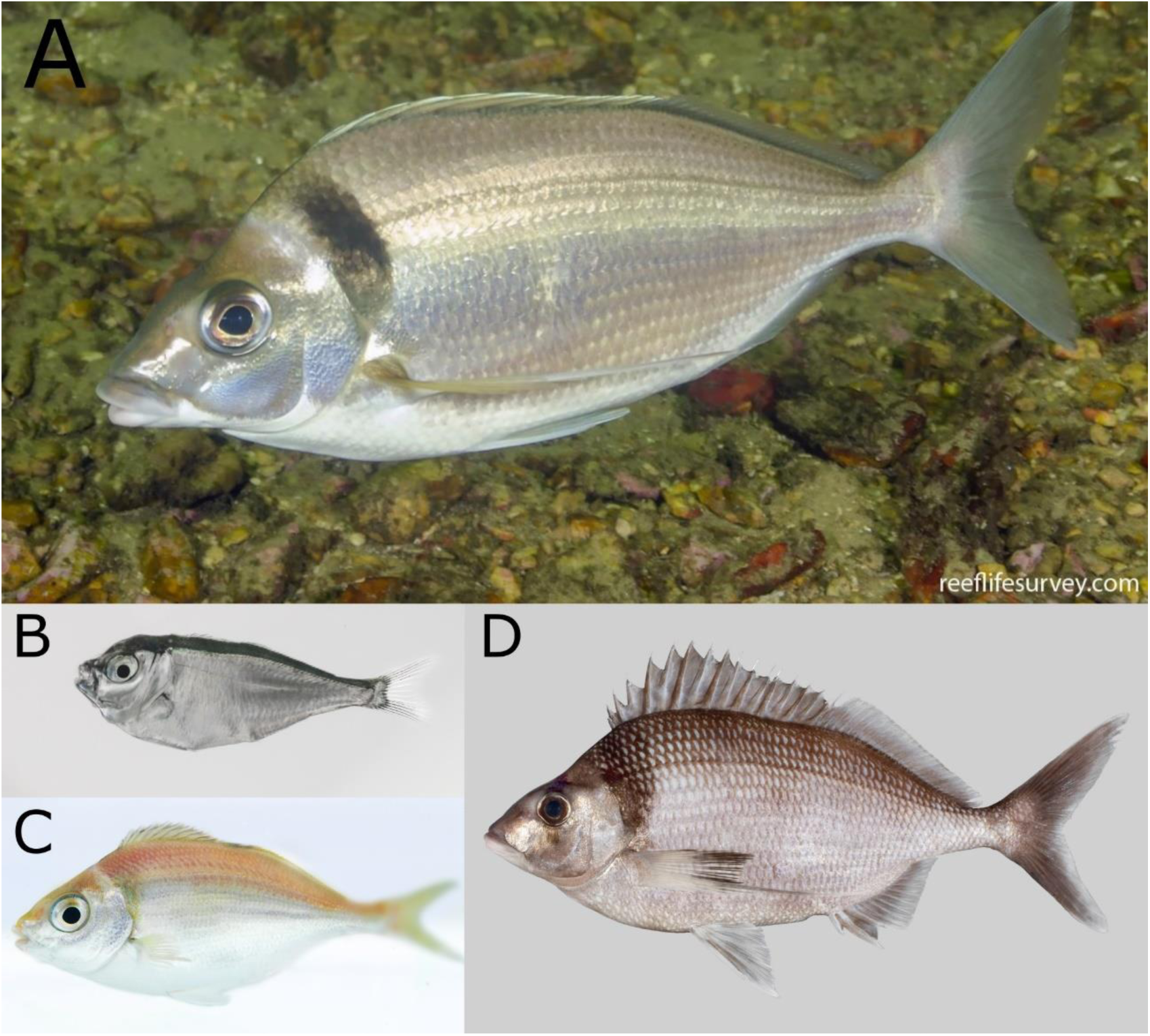
(A) An adult tarakihi (*Nemadactylus macropterus*) in Port Davey, Tasmania. In the late-larval stage (B), the individuals display a deep body with a strong ventral keel particularly adapted to pelagic life. This “paperfish” stage is thought to be an important stage for dispersal of tarakihi due to its duration (c. 10 months) and the strong influence of oceanic currents on its spatial distribution. Metamorphosis into the juvenile stage (C) occurs when the fish are 7–12 months old and 7–9 cm long. At this stage, the individuals settle in “nurseries” and become more sedentary. (D) King tarakihi (*Nemadactylus* n.sp); the main visually recognizable difference to tarakihi is the presence of a black band on the pectoral fin. Pictures by courtesy of (A) Ian Shaw, Reef Life Survey, (B-C) Robert Lamberts, the New Zealand Institute for Plant and Food Research Limited, and (D) the Museum of New Zealand Te Papa Tongarewa.

**Figure 2.**
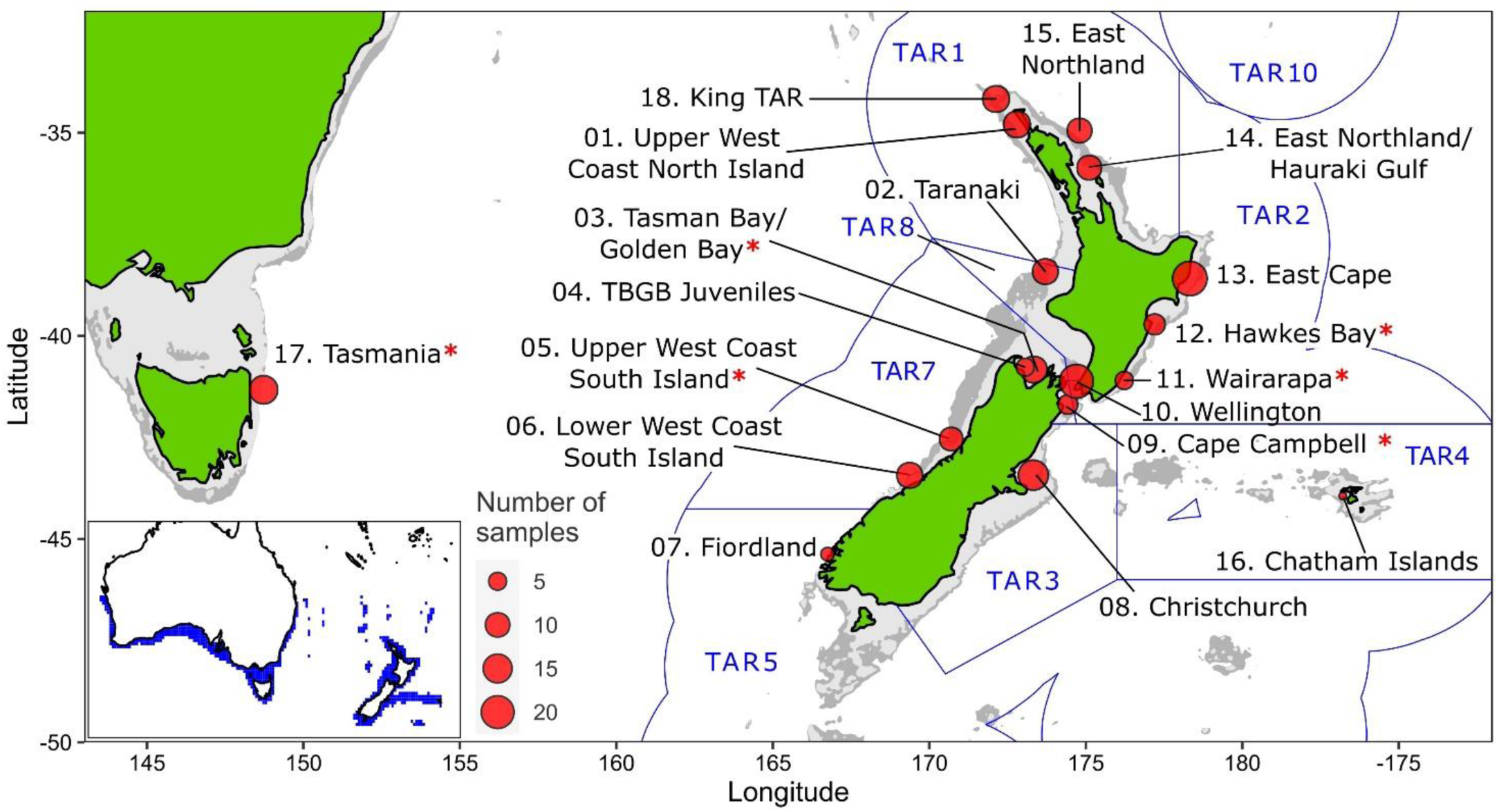
Sampling sites of tarakihi (sites 1–17) and king tarakihi (site 18) used in this study. King TAR: king tarakihi from Three Kings Islands. Red asterisks (*) indicate populations that are adaptively divergent based on outlier analysis. Blue lines and labels correspond to the New Zealand Exclusive Economic Zone and the quota management areas for tarakihi (TAR 1–5, 7–8, 10). The bathymetric contour is shown in light grey (125 m) and dark grey (1000 m). Bottom left: range of tarakihi in blue area (Kaschner et al., 2019).

DNA-based markers are particularly appropriate to provide evidence for stock boundaries, especially if a population has experienced long-term (total or partial) reproductive isolation (Waples et al., 2008; Ovenden et al., 2015; Cadrin, 2020). Six population genetic studies have been conducted on tarakihi in an effort to resolve its stock structure. Four of these studies investigated samples from around Australia and included only one location from New Zealand, using allozyme electrophoresis (Richardson, 1982; Elliott and Ward, 1994), mitochondrial DNA restriction fragment length polymorphism (Grewe et al., 1994), and microsatellite DNA markers (Burridge and Smolenski, 2003). No significant genetic structure among Australian stocks was detected. Weak but significant genetic divergence was found between New Zealand and Australia (Richardson, 1982; Elliott and Ward, 1994; Grewe et al., 1994), except in one study (Burridge and Smolenski, 2003). Two genetic studies have been conducted among New Zealand locations. The first study (Gauldie and Johnston, 1980) was conducted on c. 3,000 samples from around New Zealand. Significant population genetic differentiation was observed based on allelic frequencies at the phosphoglucomutase allozyme locus, which led to the proposition of eight geographical stock boundaries around the North and South Island (see Gauldie & Johnston (1980) for more details on these putative stock boundaries). However, it was not possible to rule out a deviation from selective neutrality for that locus. Indeed, the authors emphasized that the observed differences in allele frequencies might not be indicative of differentiation due to reproductively separated stocks, but were most probably due to an adaptive cline related to water temperature. The second study used direct sequencing of the mitochondrial DNA control region from 370 tarakihi collected around New Zealand main islands and Chatham Islands (Papa et al., 2021a). While again no overall genetic structure was found among New Zealand and Chatham Island populations, two weak genetic disjunctions were detected. The first was between the west and east coasts of the South Island, which would be consistent with biological observations based on age and size structure (Langley, 2018). The second was between Hawke’s Bay and East Northland (but not the locations between them), which may be indicative of a complex migration pattern along the east coast of North Island.

The overall goal of this study was to determine the population genetic structure of tarakihi sampled from sites around New Zealand and analysed using whole-genome resequencing. This high-resolution dataset was used to 1) characterize neutral and 2) adaptive genetic variation across sampling locations, and 3) evaluate the relationship between genetic differentiation and environmental factors. The results are compared to the current fishery stock hypotheses for tarakihi.

## 2 Materials and Methods

### 2.1 Sampling and DNA extraction

One hundred eighty-eight samples were used, including 161 tarakihi from New Zealand, 14 tarakihi from Tasmania and 12 king tarakihi (*Nemadactylus* n.sp.) from the north of New Zealand (Figure 2, Table 1). The king tarakihi phenotype is similar to tarakihi and is managed as part of the same fisheries (Figure 1D). Samples from king tarakihi were included to compare the observed levels of diversity with a close taxon. An additional specimen, caught in a fishing competition in Gisborne (East Cape), was visually identified as a king tarakihi and was added to the dataset (referred to as “GBK” for Gisborne king tarakihi). All 188 samples were sourced from specimens captured during two sampling phases. The first sampling phase took place between October 2017 and April 2018 and aimed at collecting specimens from all around New Zealand for a previous study on the population genetics of tarakihi based on a mitochondrial marker (Papa et al., 2021a). All specimens were sourced from commercial fishing companies (which in some cases conducted specific tows targeting tarakihi) with the exception of Fiordland samples that were provided by recreational fishers. Although most of the tissues collected during the first sampling phase were suitable for Sanger sequencing as applied in the mitochondrial study, DNA in these samples was generally too degraded for Illumina library preparation and whole-genome sequencing. As a result, 46 samples from phase 1 that passed the quality controls criteria were used in this study. The remaining 142 samples came from specimens captured during a second sampling phase from January 2019 to June 2020. This included fish captured by commercial fishing companies and/or as part of a broader monitoring campaign from the National Institute of Water and Atmospheric Research (NIWA), as well as specimens captured during recreational fishing competitions. An additional 14 specimens captured off Tasmania were purchased at a Sydney market through the New South Wales Department of Primary Industries program.

**Table 1.**
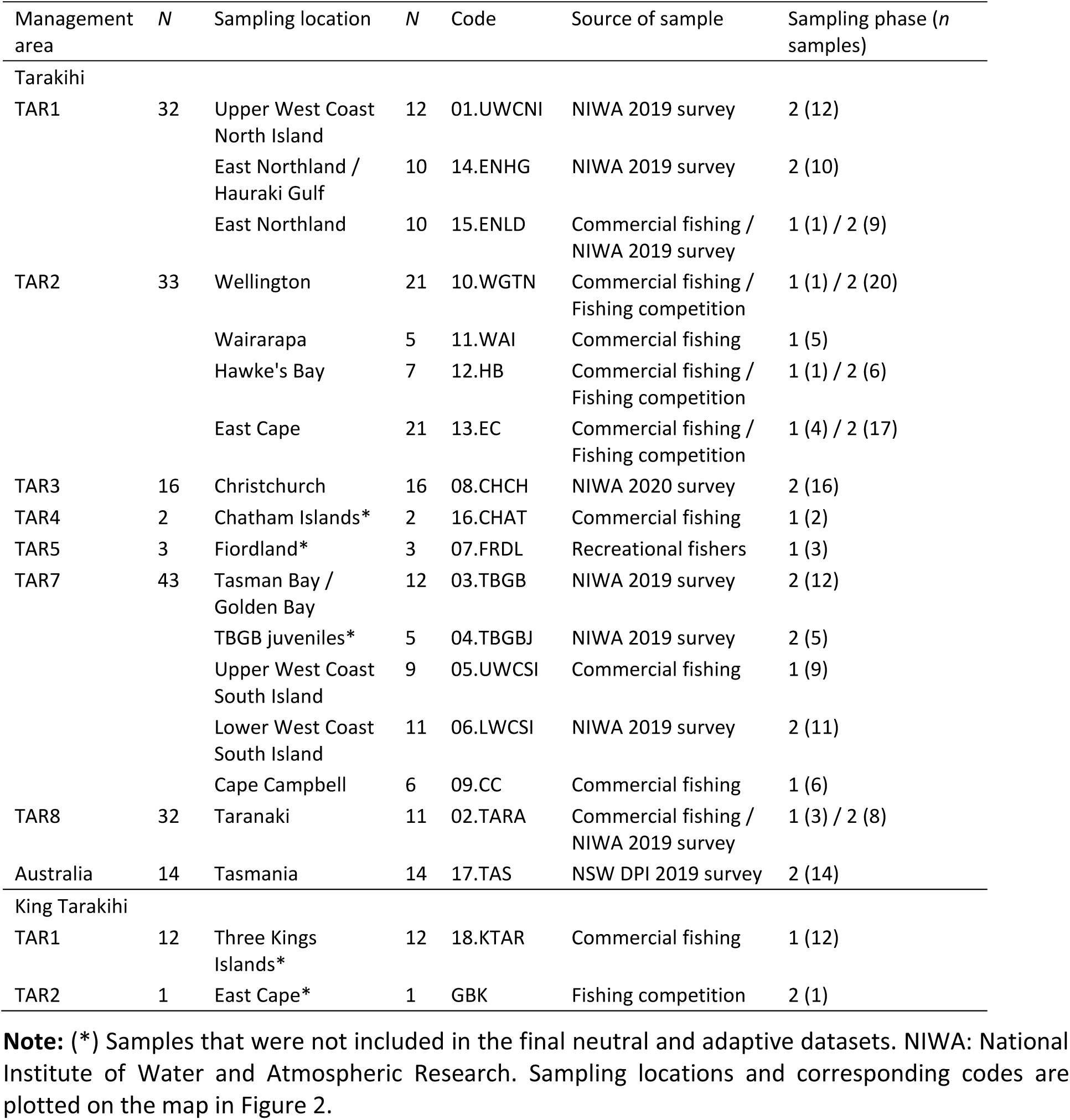
Sampling sites and sample information.

Tail muscle (phase 1) or pectoral fin (phase 2) tissue was collected from the specimens and immersed in 99% ethanol (phase 1 and 2) or DESS solution (20% DMSO, 0.25 M EDTA, NaCl saturated) (phase 2) and then stored at −20 °C. DESS was found to be more suitable to preserve DNA when tissue is sampled in the field (Oosting et al., 2020), which is generally the case when dealing with wild fisheries specimens. Total genomic DNA was extracted with a rapid salt-extraction protocol (Aljanabi and Martinez, 1997) that included an RNase step, suspended in 30–150 µl Tris-EDTA buffer (10 mM Tris-HCl pH 8.0, 0.1 mM EDTA), and stored at −20 °C. The quantity of Tris-EDTA was chosen based on visual assessment of the size of the DNA pellet. A low concentration of EDTA (0.1 mM) was used to reduce the risks of enzymatic inhibitions during library preparation. The integrity of extracted DNA samples was assessed by visualising the presence of high molecular weight DNA with agarose gel electrophoreses. Electrophoreses were run in TBE-buffered 1% agarose gels at 90 V and 400 mA for 30 min. Samples were visually classified as high, medium, or low weight depending on the intensity of DNA > 20 Kb as measured with a Lambda *Hind*III DNA ladder. Purity of DNA samples was measured with a NanoPhotometer® NP80 (Implen). Readings of 260/280 were classified as good (1.8–1.95), medium (1.6–2) or bad (any other value). Similarly, 260/230 values were classified as good (≥ 2), medium (≥ 1.4), or bad. Quantity of double-strand DNA was measured using a Qubit™ dsDNA BR Assay Kit. Samples were classified based on dsDNA concentration in a final volume of 100 µl as follows: high (≥ 50 ng/µl), medium (≥ 5 ng/µl), or low (< 5 ng/µl). DNA samples were selected based on both quality and geographic representation for DNA library preparation and sequencing.

### 2.2 Whole-Genome Sequencing

A total of 188 DNA samples were selected for whole-genome sequencing. An effort was made to sequence at least 10 specimens per sampling location, however, this could not be achieved for seven out of 18 locations. In particular, only a few DNA samples of sufficient quality could be obtained for remote locations that were sampled only in phase 1 (Chatham Islands and Fiordland). Nevertheless, a good overall representation of tarakihi fishing areas around mainland New Zealand was obtained (Figure 2), especially in the most fished management areas (TAR1, TAR2, and TAR3). DNA samples were diluted to an equal volume of 80 µl with a DNA concentration of 30 ng/µl (sometimes lower when not possible) and sent to the Australian Genome Research Facility (AGRF, Melbourne, Australia) for DNA library preparation and sequencing. The Illumina DNA shotgun library was prepared following the Nextera DNA FLEX low volume protocol with Nextera DNA Combinatorial Dual Indexes (Illumina) for insert sizes 300–350 bp. Sequencing of 150 bp paired-end reads was performed on NovaSeq 6000 (Illumina) with NovaSeq 6000 S4 Reagent Kit and NovaSeq XP 4-Lane Kit for 300 cycles. Each lane contained 96 wells, and each individual was sequenced in one well. Since the sequencing yield of each lane was 700–800 Gb, it was expected that each individual would be sequenced for c. 8Gb. The genome size was estimated to be around 700 Mb based on the C-value of 0.72 for *Cheilodactylus fuscus* on the Animal Genome Size Database (http://www.genomesize.com). The sequencing coverage was thus estimated to be c. 11× per sample. Base calling, quality scoring, and de-multiplexing were performed by sequencing provider with RTA3 software v3.3.3 and Illumina bcl2fastq pipeline v2.20.0.422.

### 2.3 Quality control and pre-processing

The quality of the paired-end reads was assessed with FastQC v0.11.7 (Andrews, 2018) and MultiQC v1.7 (Ewels et al., 2016) before and after trimming. Raw reads were trimmed for adapter contamination and unpaired reads were filtered out using Trimmomatic v0.39 (Bolger et al., 2014) with the following parameters: paired-end mode, phred 33 base quality encoding, maximum allowed mismatch count = 2, match accuracy between adapter-ligated reads = 30, match accuracy between adapter and read = 10. Nextera adapters were targeted using the file `NexteraPE-PE.fa` provided by Trimmomatic.

### 2.4 Genotyping

Trimmed and filtered paired reads were mapped to the tarakihi reference genome (1,214 scaffolds) assembled in a previous study (Papa et al., 2021c) by using the Burrows-Wheeler Alignment (BWA) method with bwa-kit v0.7.15 (Li and Durbin, 2009). In brief, a BWA index was created for the reference genome, as well as a SAM index with SAMtools v1.9 (Li et al., 2009). For each specimen, the forward and reverse reads were mapped to the genome with the command `bwa mem –a –M` to flag all single-end and unpaired reads and mark shorter split hits as secondary. Duplicated reads were marked using Picard v2.18.20 (Broad Institute, 2019) `MarkDuplicates` with default settings. Genotype likelihoods were produced from the BAM files using bcftools v1.9 (Li, 2011) command `mpileup.` The minimum mapping quality was set to 10, the mapping quality downgrading coefficient was set to 50 for reads with excessive mismatches, and only reads mapped in proper pair were kept (e.g. no secondary alignment or duplicates). SNPs were then called with bcftools multi-allelic caller on default parameters, outputting only variant sites and skipping indels. Genotype likelihoods and SNPs calling were both performed separately on each of the 1,214 assembly scaffolds and results were then merged into a genome-wide SNP dataset using bcftools `concat`. The final BCF file was sorted and indexed after having renamed the headers.

### 2.5 Variant filtering

Several SNP datasets were produced and analysed separately in this study (Figure 3). The first was a pruned dataset that contained the 188 tarakihi and king tarakihi individuals, where SNPs were filtered for minimum quality criteria and pruned for linkage and Hardy-Weinberg disequilibrium. The second was a dataset of neutral SNPs (hereafter referred to as the “neutral dataset”) that only included the 250 longest scaffolds from locations with ≥ 5 sampled adult tarakihi individuals (king tarakihi, Fiordland, Chatham Islands, Tasman Bay/Golden Bay (TBGB) juveniles, and GBK were discarded). SNP filtering for the neutral dataset was identical to the pruned dataset but with an additional step of filtering out potentially adaptive outliers (see below). The 250 longest scaffolds were retained because (1) they contained more than 90% of the total number of bases in the tarakihi reference genome (L90 = 219 (Papa et al., 2021c)), (2) for computational efficiency, and (3) because the default parameters used for the OutFLANK analysis (see below) were not optimal anymore to fit the *F*_ST_ curve in some scaffolds past that number, which means they would have had to be manually tuned for the c. 1,000 remaining scaffolds. Chatham Islands, Fiordland and the juvenile TBGB samples were discarded at that stage because of their low number of samples (two, three, and five, respectively), which was considered too low to confidently predict population allele frequencies. The GBK specimen was discarded because of its dubious field identification, and the king tarakihi were discarded for being a different species with a different demographic history. The third and fourth datasets were outlier-based, presumably adaptive datasets containing the same individuals and scaffolds as the neutral dataset, but including only strong candidates for adaptive loci (see below). The fifth, environment-adaptive, dataset was obtained through locus–environment association analysis (see below). Filtering of variant sites was performed with VCFtools v0.1.16 (Danecek et al., 2011). Filtering steps were primarily carried out separately on each scaffold, and resulting VCF files were merged with Picard v2.18.20.

**Figure 3.**
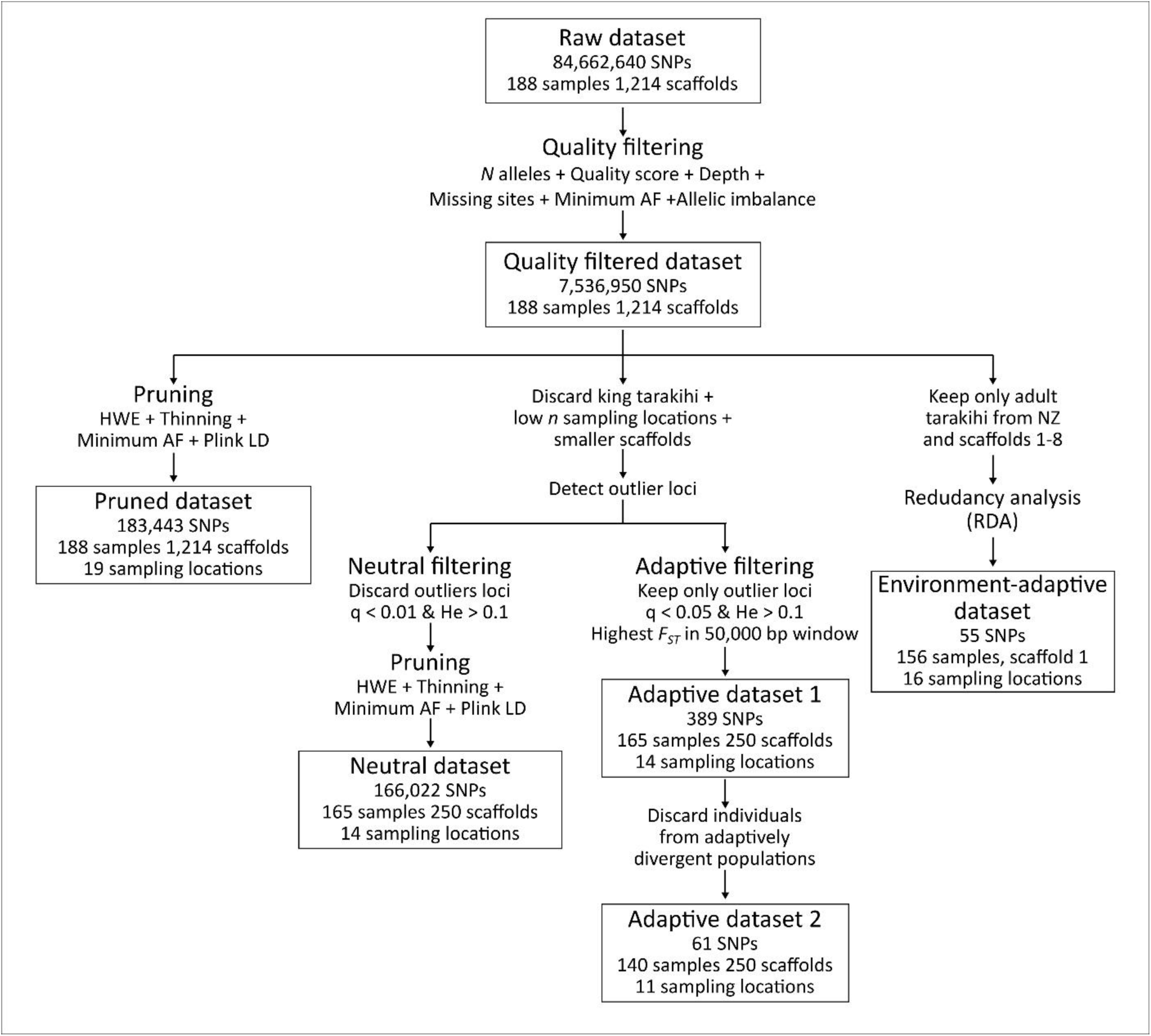
Variant filtering pipeline applied on the raw SNP dataset, resulting in one intermediary dataset (quality-filtered) and five final datasets (pruned, neutral, adaptive 1, adaptive 2, and environment-adaptive).

#### Quality filtering

Only bi-allelic sites with a minimum quality of 600 were retained (`--max-alleles 2 --min-alleles 2 --minQ 600`). The minimum allelic depth for sites in an individual was set to three and the mean depth of sites across all individuals was set between eight and twenty-five (-`-minDP 3 --min-meanDP 8 --max-meanDP 25`). Sites that were missing in more than 5% of individuals and sites with a minor allele frequency lower than 1% were filtered out (`--max-missing 0.95 --maf 0.01`). For each site, potential allelic imbalance was detected by running a binomial test on the sum of all reference alleles and the sum of all alternative alleles across all sequenced reads of heterozygous individuals with a custom R script, using the genotype (GT) and allelic depth (AD) format information. Sites were filtered out with VCFtools `(--exclude-positions`) if the total proportion of reference and alternative alleles was significantly different from 50% (*P* ≤ 0.05 after correction for false discovery rate) according to the binomial test.

#### Linkage disequilibrium

In order to choose a threshold value for thinning in the pruning step, linkage disequilibrium decay was plotted for the 30 longest scaffolds on the quality-filtered dataset. For this, the squared correlation coefficients between genotypes of sites separated by a maximum of 50,000 bp were calculated with VCFtools (`--geno-r2 --ld-window-bp 50000`). Nucleotide position and *R*^2^ values for a random subset of a million sites in each scaffold were then plotted with a custom R script (Oosting, 2021). The analysis was run twice, once on the 188 individuals, and once without including king tarakihi specimens.

#### Pruning

Sites that were significantly deviating from Hardy-Weinberg equilibrium were filtered out, as well as sites occurring within a distance of 1,500 bp from one another in the scaffolds (`--hwe 0.05 --thin 1500`). A minimum allele frequency of 0.01 was applied a second time. The presence of remaining linkage disequilibrium after thinning was detected with PLINK v1.90 (Chang et al., 2015), using a window size of 50 sites, shifting every five sites and using a pairwise *R*^2^ threshold of 0.2 (`--indep-pairwise 50 5 0.2`). These SNPs were then filtered out with VCFtools.

#### Neutral filtering

Sites that were potentially under selection were detected with OutFLANK v0.2 (Whitlock and Lotterhos, 2015) using default parameters. Loci flagged as outliers with a minimum heterozygosity of 0.1 (`He > 0.1`) and a false discovery threshold below 1% (`qvalues < 0.01`), were filtered out of the dataset with VCFtools.

#### Adaptive filtering

In parallel with the neutral filtering, OutFLANK v0.2 was used on the pruned dataset to flag outlier loci with a minimum heterozygosity of 0.1 and a false discovery threshold (q-value) below 0.05. This analysis was run twice: the sampling locations that were detected as differentiated were discarded between the first and second analyses to detect SNPs with finer population structure. After each analysis, a custom R script adapted from Oosting (2021) was used to select only the site with the highest *F*_ST_ in each non-overlapping 50,000 bp sliding window in order to obtain a set of independent, presumably adaptive SNPs.

### 2.6 SNP data analysis

Number of reads per individuals and mean GC content were reported with FastQC v0.11.7 and MultiQC v1.7. Mean read depth of variant sites and proportion of missing sites in each individual were calculated with VCFtools v0.1.16. Observed heterozygosity and number of fixed alleles were obtained with dartR v1.9.6 (Gruber et al., 2018). Analyses of Molecular Variance (AMOVA) were conducted on the neutral dataset with poppr v2.9.2 (Kamvar et al., 2014) using hierarchical models of several groupings based on management areas, New Zealand coasts, and sampling locations. Associated *p*-values were computed with ade4 v1.7.16 (Dray and Dufour, 2007) function `randtest` using 999 replicates. Principal component analyses (PCA) were run on the pruned, neutral, and adaptive SNP datasets with dartR v1.9.6 using default parameters and results were projected onto axes with ggplot2 v3.3.3 (Wickham, 2009). Successive K-means cluster identification and discriminant analyses of principal components (DAPC) were performed with adegenet v2.1.3 (Jombart, 2008). See parameters used in Supplementary Table 1.

Pairwise weighted *F*_ST_ values (Weir and Cockerham, 1984) were computed with StAMPP v1.6.1 (Pembleton et al., 2013) on the neutral and adaptive datasets. Associated *p*-values and 95% confidence intervals were generated by running 1,000 bootstraps across loci and a false discovery rate correction (Benjamini and Hochberg, 1995) was applied to the *p*-values. The pairwise weighted *F*_ST_ values were plotted onto heat maps clustered via dendrogram using pheatmap 1.0.12 (Kolde, 2019). Population structure was tentatively inferred in the neutral dataset with fastSTRUCTURE v1.0.3 (Raj et al., 2014). The number of populations (*K*) was tested from 1 to 10, and the value of *K* that best explained the observed structure was chosen with the utility tool `chooseK.py`.

A test of isolation by distance (IBD) was performed using a Mantel test (999 replicates) with ade4 v1.7.16 on the neutral SNP dataset. The analysis was restricted to the samples from New Zealand locations. The geographic distances between sample coordinates were estimated with gdistance v1.3.6 (van Etten, 2017) by applying a ‘least-cost distance’ model of geographic dispersal where travel is restricted to the ocean. For this, a shapefile of New Zealand was rasterized with raster v3.4.5 (Hijmans, 2019), and each cell grid corresponding to the ocean and the land were assigned movement costs of 0 and 1, respectively. Sampling location coordinates were obtained by calculating the mean between the start and stop trawl latitudes and longitudes provided by the fishing vessels, or approximated arbitrarily when not provided. Pairwise weighted *F*_ST_ values between sampling locations were used for the matrix of genetic distances.

### 2.7 Genotype-environment association analysis

Environmental feature data was obtained with the R package sdmpredictors v0.2.9 (Bosch, 2020). The complete dataset available from Bio-ORACLE v1, v2.1, and MARSPEC (Tyberghein et al., 2012; Sbrocco and Barber, 2013; Assis et al., 2018) was compiled into a total environmental dataset of 362 variables. Since the locus-environment association method used is a regression-based analysis, it was necessary to check for collinearity (i.e. non-independence) between the environmental variables before proceeding (Dormann et al., 2013). This was done in several steps. First, the 362 variables were manually clustered into groups that each measured the same physical, chemical, or biological process (Supplementary Table 2). As an example, the group “dissolved oxygen”, included 25 variables, such as sea surface mean concentration, minimum concentration at mean depth, or concentration range at the bottom. For each of these groups, the R package psych v2.1.3 (Revelle, 2021) was used to assess the correlation level among variables. A set of weakly correlated variables (Pearson correlation |*R*^2^| < 0.7) was manually selected from each group and compiled in a second dataset of 48 variables (Supplementary Figure 1). The Pearson correlation and associated significance were computed among these 48 variables with the R package Hmisc v4.5-0 (Harrell, 2021). All variables that were significantly correlated to “mean temperature at mean depth” (*P* ≤ 0.05) were discarded. Temperature was chosen because it is usually correlated with many other environmental variables as it is associated with, or drives, many physical, chemical, and biological processes, and the mean value at mean depth was chosen to reflect the biology of tarakihi, which is a demersal, inshore species. Variables that were considered to be not biologically relevant for the studied species (cloud fraction, ice cover, East/West aspect and North/South aspect) were also discarded. This led to a third dataset of nine variables (Supplementary Figure 2). Three variables (profile curvature, plan curvature, and mean chlorophyll concentration at minimum depth) were further discarded for being correlated to concavity and mean primary production at the bottom (|*R*^2^| > 0.5, *P* < 0.001). This reduced the dataset to a final six lowly correlated variables: salinity range at mean depth, mean primary production at minimum depth, concavity (which is indicative of the location being on a slope or in a valley), mean temperature at mean depth, mean diffuse attenuation coefficient (indicator of water clarity), and mean iron concentration at the sea surface (Supplementary Figure 2).

Association between genotypes and environmental variables was assessed using a Redundancy Analysis (RDA). RDA is a multivariate, ordination-based locus-environment association method that is effective at detecting local adaptation on multiple loci under numerous demographical, biological, and sampling scenarios (Rellstab et al., 2015; Forester et al., 2016, 2018). The analysis was performed separately on the eight longest scaffolds using the quality-filtered SNP dataset (Figure 3), without including king tarakihi, GBK, specimens from Tasmania, and juveniles from TBGB. Missing genotype data was estimated using the most common genotype at each locus. The RDA was run with the package vegan v2.5.7 (Oksanen et al., 2020) on the genotype data and the six environmental variables cited above, with parameter scale = TRUE to scale observations to unit variance. The significance of each RDA model, as well as each constrained axis, was assessed with vegan v2.5.7 function `anova.cca` using 999 permutations. Loci that were strong candidates for local adaptation were selected by identifying the SNPs with scores ± 3.5 standard deviations from the mean score of each significant constrained axis. The position in the genome and putative associated function of the selected loci were obtained using the tarakihi reference genome assembly annotation produced in Papa et al. (2021c).

### 2.8 General bioinformatics tools

All SAM files were converted to sorted BAM files with SAMtools `sort` and BAM files were indexed with SAMtools `index`. Alignments statistics were computed with SAMtools `flagstats` and BamTools v2.5.1 (Barnett et al., 2011) `stats`. VCF files were imported and converted to genlight objects in R with vcfR v1.12.0 (Knaus and Grünwald, 2017). dartR v1.9.6 was used for manipulations of genlight objects (e.g. population assignment, filtering of individuals and monomorphic loci). Analyses were performed on Rāpoi, the Victoria University of Wellington high-performance computer cluster. Analyses requiring R scripts were performed in R v4.02 (R Core Team, 2020) on RStudio (RStudio Team, 2020).

## 3 Results

### 3.1 Sequencing

Following quality filtering and adapter trimming, a total of 12.1 billion Illumina reads, with an average length of 150 bp, were obtained from 188 individuals. The mean GC content was 44.8%. All individual samples passed all the FastQC criteria. The number of reads per sample ranged from 17.1 million to 124.0 million (64.1 million on average) (Supplementary Figure 3A). This translated to a mean read sequencing depth per individual of 16.9×, ranging from 4.5× to 32.7×, with only eight individuals below 8×.

### 3.2 SNP datasets

Quality filtering of variants led to a total of 7,536,950 high-quality bi-allelic SNPs with a mean depth per individual of 12.7× (Supplementary Figure 3B). The “pruned” dataset that included all scaffolds and individuals was made up of 183,443 high-quality, independently segregating SNPs. To obtain the neutral dataset, 23 individuals were discarded and the analysis was restricted to the 250 longest scaffolds (see Methods). This final neutral dataset contained 166,022 high-quality, independently segregating, neutrally evolving bi-allelic SNPs (Figure 3). The mean SNP depth per individual in that dataset was 12.0× (0.04x-23×) and the number of missing sites per individual ranged from 5 to 164,506, with 151 individuals (92% of total) missing less than 400 sites (Supplementary Figure 3C).

There was a clear difference in observed heterozygosity between tarakihi and king tarakihi specimens in the quality-filtered dataset, with an average of 0.12 and 0.06, respectively (Supplementary Figure 3B). The GBK specimen had a heterozygosity level typical of a tarakihi. In the neutral dataset, the mean observed heterozygosity was 0.13, ranging from 0.09 to 0.15. The levels of heterozygosity did not significantly vary among locations but were directly related to the mean SNP depth, which in turn was correlated to the number of missing sites per individual. The relationship between heterozygosity levels and depth of coverage was especially evident in the 14 samples with a heterozygosity < 0.12, which all had a mean depth < 7x.

In the high-quality total SNP dataset, 84,144 allelic differences were fixed between the king tarakihi and the tarakihi specimens (not including GBK). In comparison, the average number of fixed mutations among all tarakihi locations ranged from 1 (East Cape) to 145 (Chatham Islands), with only TBGB juveniles, Wairarapa, Tasmania, Fiordland, and Chatham Islands having more than 20. There was only one fixed mutation between GBK and the tarakihi specimens.

### 3.3 Linkage Disequilibrium

Linkage disequilibrium in tarakihi was low overall, with mean pairwise *R*^2^ values never exceeding 0.2, even between nucleotide sites less than 100 bp apart (Figure 4). Linkage disequilibrium decay was also rapid: both *R*^2^ and mean *R*^2^ values plotted on the distance between nucleotides always reached a plateau between 500 and 1,500 bp (Figure 4). The threshold limit for the thinning step in the variant filtering pipeline was thus set to a conservative 1,500 bp. Interestingly, when the same analysis was run while including the king tarakihi specimen, the *R*^2^ values were generally higher and more uniform along the length of the scaffolds (Supplementary Figure 4).

**Figure 4.**
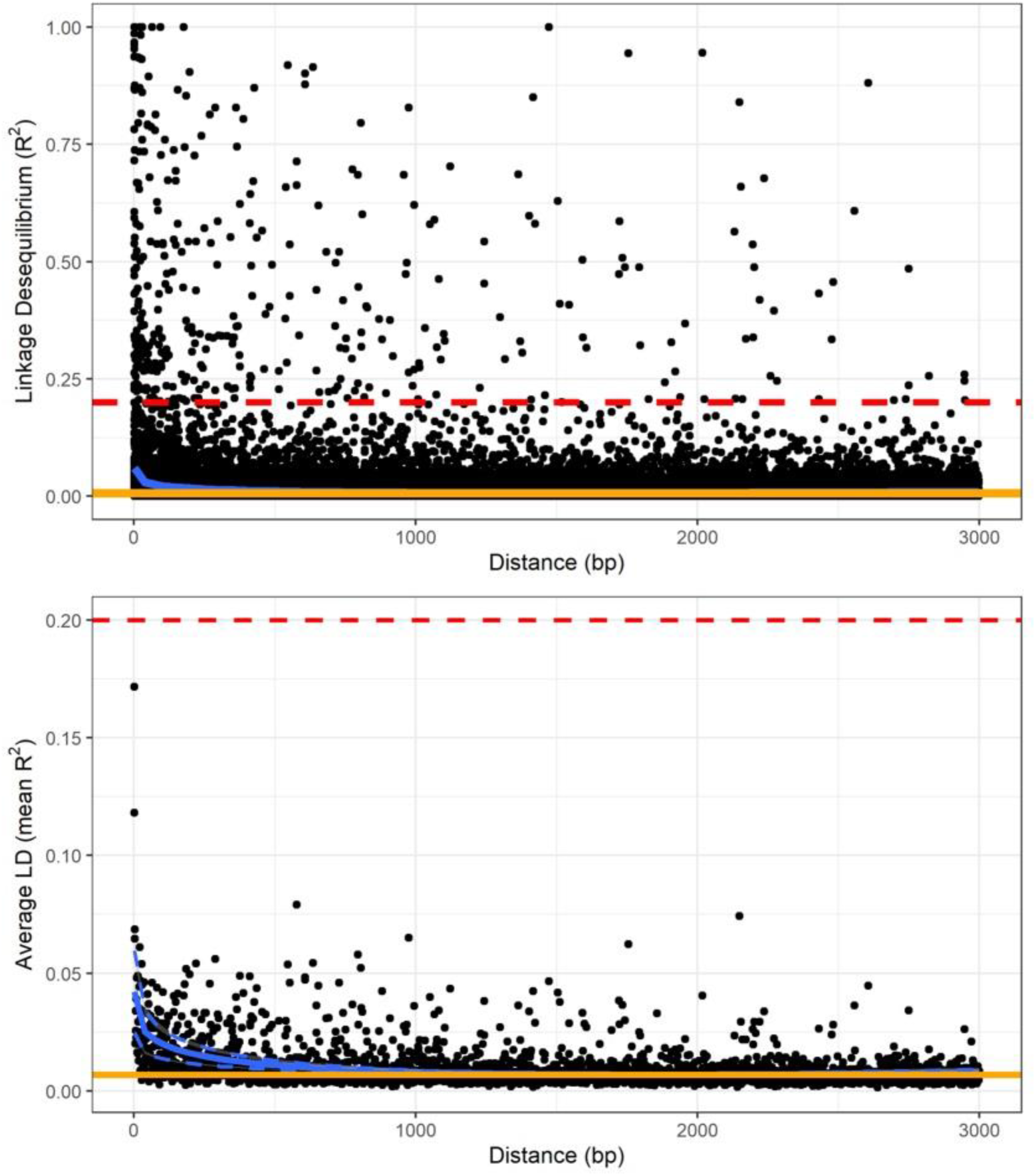
Linkage disequilibrium decay over genetic distance on scaffold 1, calculated on the quality-filtered SNP dataset, minus king tarakihi specimens. The horizontal red dashed line shows the threshold of 0.2 that is commonly applied to identify independent degradation of nucleotide sites. The orange line is the background level of linkage disequilibrium (intercept). The blue line is the trend of linkage disequilibrium decay fitted to the plot (minimum and maximum variance in dashed lines for the mean).

### 3.4 Population structure

Several AMOVAs were run on the neutral and the adaptive datasets to assess the proportion of genetic variance that could be explained by pre-assigned groupings in these datasets. Groupings of sampling locations included management areas (with or without including Tasmania) and New Zealand coasts (i.e. two groups: west coast and east coast). Overall, there was no significant genetic structure among management areas, coasts, or sampling locations when analysing the neutral dataset (Table 2). The only significant genetic differentiation (Ф = 0.001, *P* = 0.001), was detected among sample locations without any *a priori* broader grouping, and only when the Tasmania location was included, which means the differentiation between New Zealand and Tasmania is driving that result. Conversely, the AMOVAs based on presumably adaptive SNPs showed sample locations to always be significantly differentiated (*P* ≤ 0.01) with the percentage of explained variation ranging from 6.1% to 20.5% depending on the grouping used (Table 2), while the majority of the variation was still within individuals (76.5%–93.32%, *P* ≤ 0.01). No significant genetic structure was detected among management areas or between the west and east coasts of New Zealand in the adaptive dataset.

**Table 2.** Results from analysis of molecular variance performed on the neutral and adaptive SNP datasets, with seven *a priori* groupings.

| *A priori* grouping | *N* ind. | *N* locs | *N* groups | Among groups |  |  | Between locs within groups |  |  | Between ind. within locs |  |  | Within individuals |  |  |
| --- | --- | --- | --- | --- | --- | --- | --- | --- | --- | --- | --- | --- | --- | --- | --- |
|  |  |  |  | %Var | Φ | *P* | %Var | Φ | *P* | %Var | Φ | *P* | %Var | Φ | *P* |
| Neutral |  |  |  |  |  |  |  |  |  |  |  |  |  |  |  |
| no grouping | 165 | 14 |  |  |  |  | 0.084 | 0.001 | **0.001** | -0.260 | -0.003 | 0.565 | 100.176 | -0.002 | 0.593 |
| no grouping, NZ only | 151 | 13 |  |  |  |  | -0.011 | 0.000 | 0.769 | -0.314 | -0.003 | 0.625 | 100.324 | -0.003 | 0.640 |
| NZ TAR areas + AU | 165 | 14 | 6 | 0.104 | 0.001 | 0.123 | -0.005 | 0.000 | 0.725 | -0.260 | -0.003 | 0.609 | 100.161 | -0.002 | 0.586 |
| NZ TAR areas only | 151 | 13 | 5 | 0.002 | 0.000 | 0.501 | -0.012 | 0.000 | 0.731 | -0.314 | -0.003 | 0.611 | 100.324 | -0.003 | 0.620 |
| NZ West vs. East | 130 | 12 | 2 | 0.002 | 0.000 | 0.261 | -0.009 | 0.000 | 0.671 | -0.322 | -0.003 | 0.592 | 100.329 | -0.003 | 0.642 |
| NZ North Island: West vs. East | 76 | 7 | 2 | -0.001 | 0.000 | 0.497 | 0.010 | 0.000 | 0.462 | 0.061 | 0.001 | 0.417 | 99.930 | 0.001 | 0.449 |
| NZ South Island: West vs. East | 54 | 5 | 2 | 0.018 | 0.000 | 0.109 | -0.033 | 0.000 | 0.967 | -0.861 | -0.009 | 0.654 | 100.876 | -0.009 | 0.678 |
| Adaptive |  |  |  |  |  |  |  |  |  |  |  |  |  |  |  |
| no grouping | 165 | 14 |  |  |  |  | 20.459 | 0.205 | **0.001** | 1.176 | 0.015 | 0.10 | 78.365 | 0.216 | **0.001** |
| no grouping, NZ only | 151 | 13 |  |  |  |  | 6.060 | 0.061 | **0.001** | 1.632 | 0.017 | 0.07 | 92.308 | 0.077 | **0.001** |
| NZ TAR areas + AU | 165 | 14 | 6 | 16.006 | 0.160 | 0.202 | 6.341 | 0.075 | **0.001** | 1.148 | 0.015 | 0.10 | 76.505 | 0.235 | **0.001** |
| NZ TAR areas only | 151 | 13 | 5 | -2.073 | -0.021 | 0.969 | 7.786 | 0.076 | **0.001** | 1.638 | 0.017 | 0.08 | 92.649 | 0.074 | **0.001** |
| NZ West vs. East | 130 | 12 | 2 | -0.634 | -0.006 | 0.970 | 6.923 | 0.069 | **0.001** | 1.609 | 0.017 | 0.08 | 92.102 | 0.079 | **0.001** |
| NZ North Island: West vs. East | 76 | 7 | 2 | -1.816 | -0.018 | 0.960 | 7.140 | 0.070 | **0.001** | 3.074 | 0.032 | 0.06 | 91.601 | 0.084 | **0.001** |
| NZ South Island: West vs. East | 54 | 5 | 2 | -1.436 | -0.014 | 0.909 | 7.181 | 0.071 | **0.001** | 0.931 | 0.010 | 0.32 | 93.323 | 0.067 | **0.001** |

**Note:** %Var = variance component in percentage of the total variation. The three ‘West vs. East’ groupings did not include the Wellington location. Significant *p*-values ( $\leq 0.05$ ) are in bold.

The PCA performed on the pruned dataset containing all 188 individuals showed that king tarakihi (18.KTAR) and the tarakihi from Tasmania (17.TAS) form two distinct clusters that are separate from all New Zealand tarakihi specimens (Figure 5). The differentiation between these two groups drove most of the variation of the first axis and the second axis, which explained 4.61% and 0.72% of the total variation, respectively. No structure was apparent among the New Zealand populations: sampling locations were randomly distributed on the third and fourth axes (Figure 5). Similarly, the DAPC conducted on *K*-means clustering did not infer any groups related to sampling locations among New Zealand tarakihi (Supplementary Figures 5 & 6). In both PCA and DAPC, the GBK specimen was always grouped within the tarakihi individuals and did not display any particular deviation from them, thus challenging its field identification as a king tarakihi. The PCA and DAPC conducted on the neutral dataset gave identical results, in that Tasmania was separated from New Zealand but no structure could be detected among New Zealand locations (Supplementary Figures 7 - 9).

**Figure 5.**
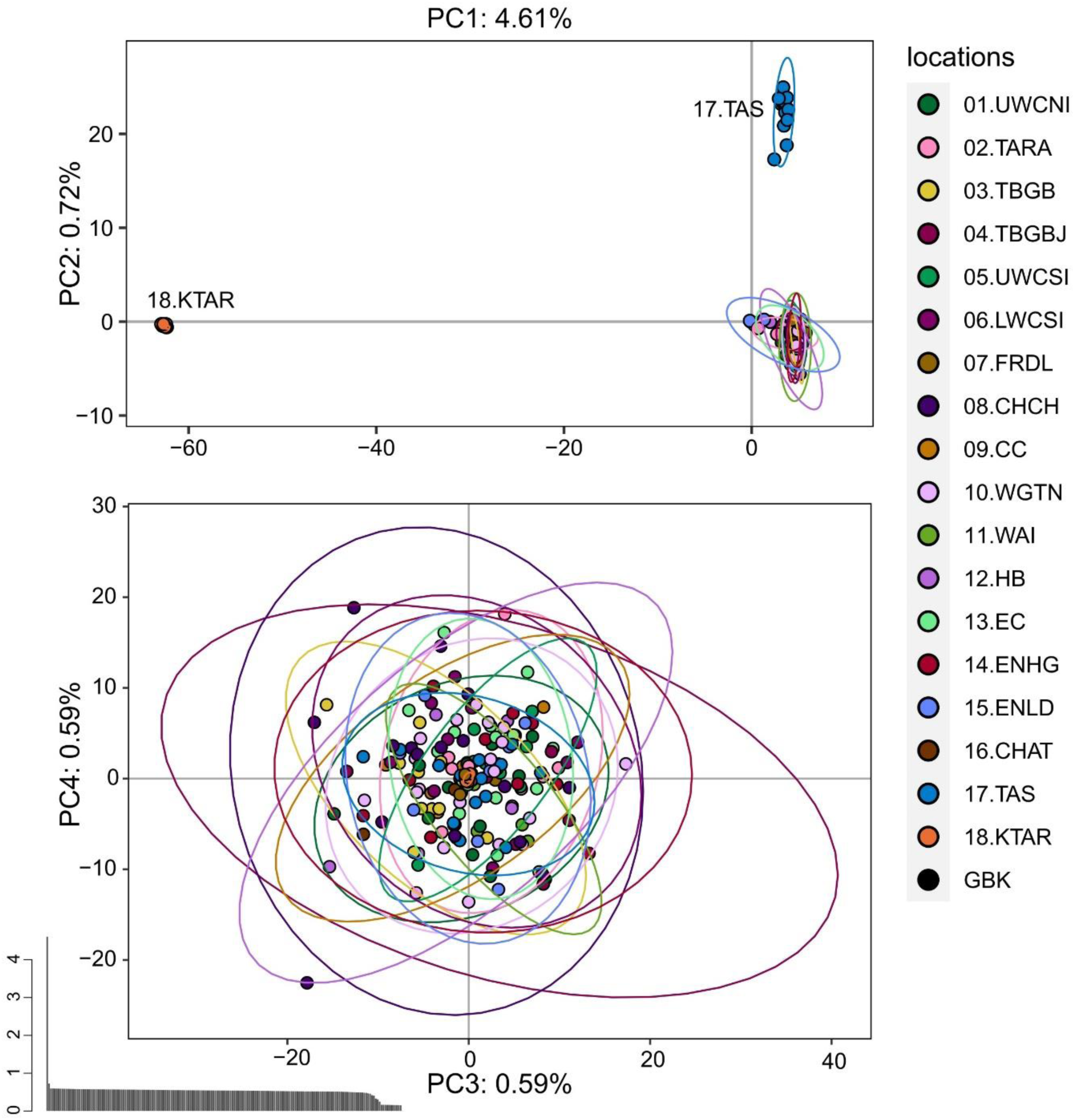
Principal component analysis of the pruned SNP dataset that includes 183,443 loci from 188 tarakihi and king tarakihi. Ellipses represent the 95% confidence intervals. Top: Axes 1 and 2. Bottom: Axes 3 and 4. Bottom left: Eigenvalues. Sampling location codes as referred to in Table 1.

PCA and DAPC performed on the adaptive dataset (389 SNPs) showed a genetic differentiation of Tasmania (17.TAS), Wairarapa (11.WAI), and Cape Campbell (09.CC) populations from the remaining tarakihi in New Zealand (Figure 6, Supplementary Figure 10). Each of these groups explained most of the variation on the first, second, and third axes of the PCA, which accounted for 25.63%, 2.41%, and 1.81% of the total variation, respectively (Supplementary Figure 10).

**Figure 6.**
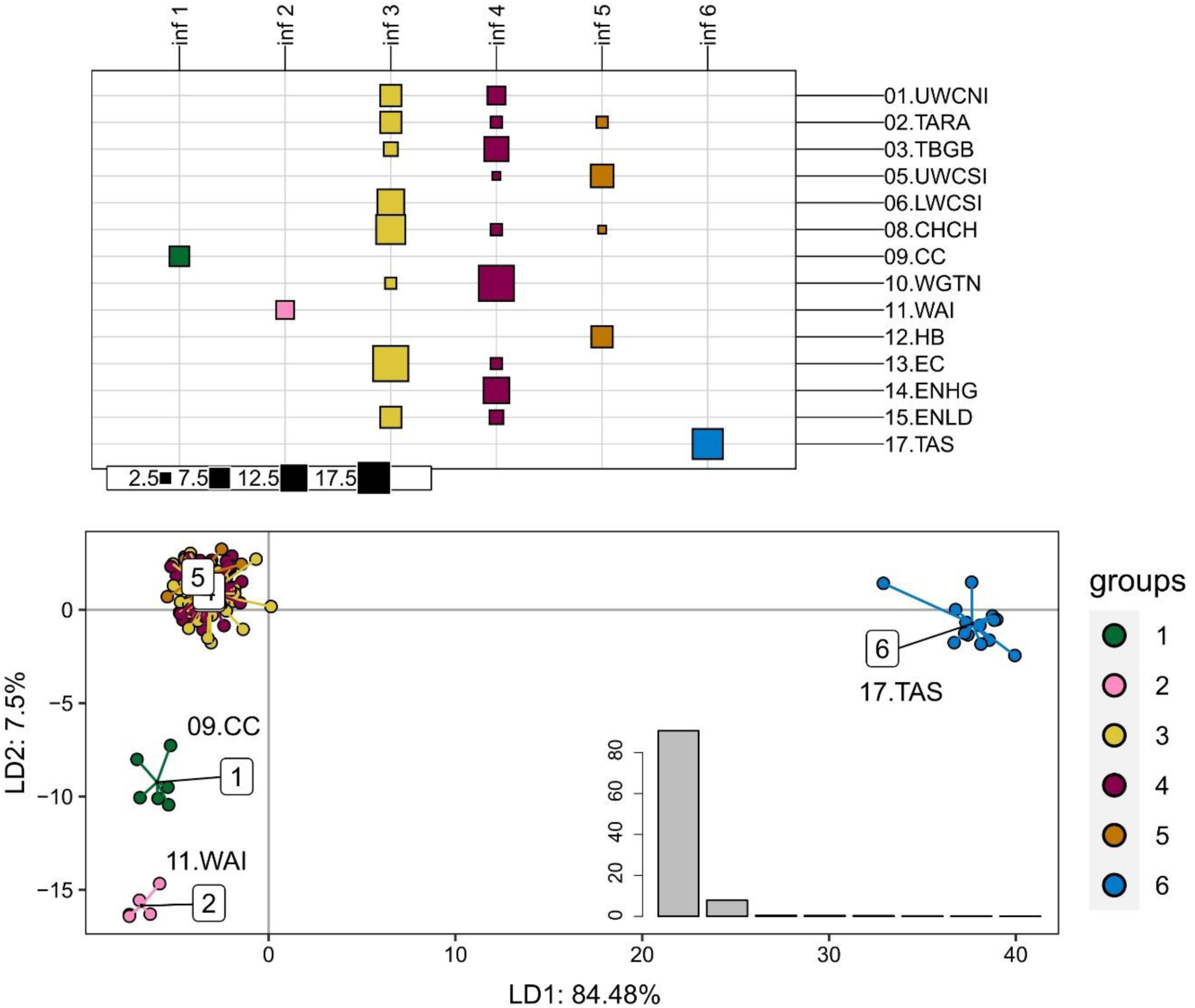
Discriminant analysis of principal components of the first adaptive SNP datasets that include 389 loci from 165 tarakihi. Top: results of the *K-*means clustering analysis, with colours corresponding to inferred groups (inf). Bottom: Projection of the DAPC based on inferred groups. Bottom right: Discriminant analysis eigenvalues. Sampling location codes as referred to in Table 1.

The PCA and DAPC analyses were performed a second time on the adaptive dataset, but this time the Tasmania, Wairarapa, and Cape Campbell locations were discarded before the outlier analysis (see Methods), which resulted in a second adaptive dataset of 61 SNPs from 140 individuals. The DAPC discriminated three additional groups that corresponded to sampling locations (Figure 7): Upper West Coast of South Island (05.UWCSI), Hawke’s Bay (12.HB), and a group from Tasman Bay/Golden Bay (03. TBGB) that also included one individual from Wellington (10.WGTN) and one from the Upper West Coast of North Island (01.UWCNI). These results were partially observable on the PCA (Supplementary Figure 11).

**Figure 7.**
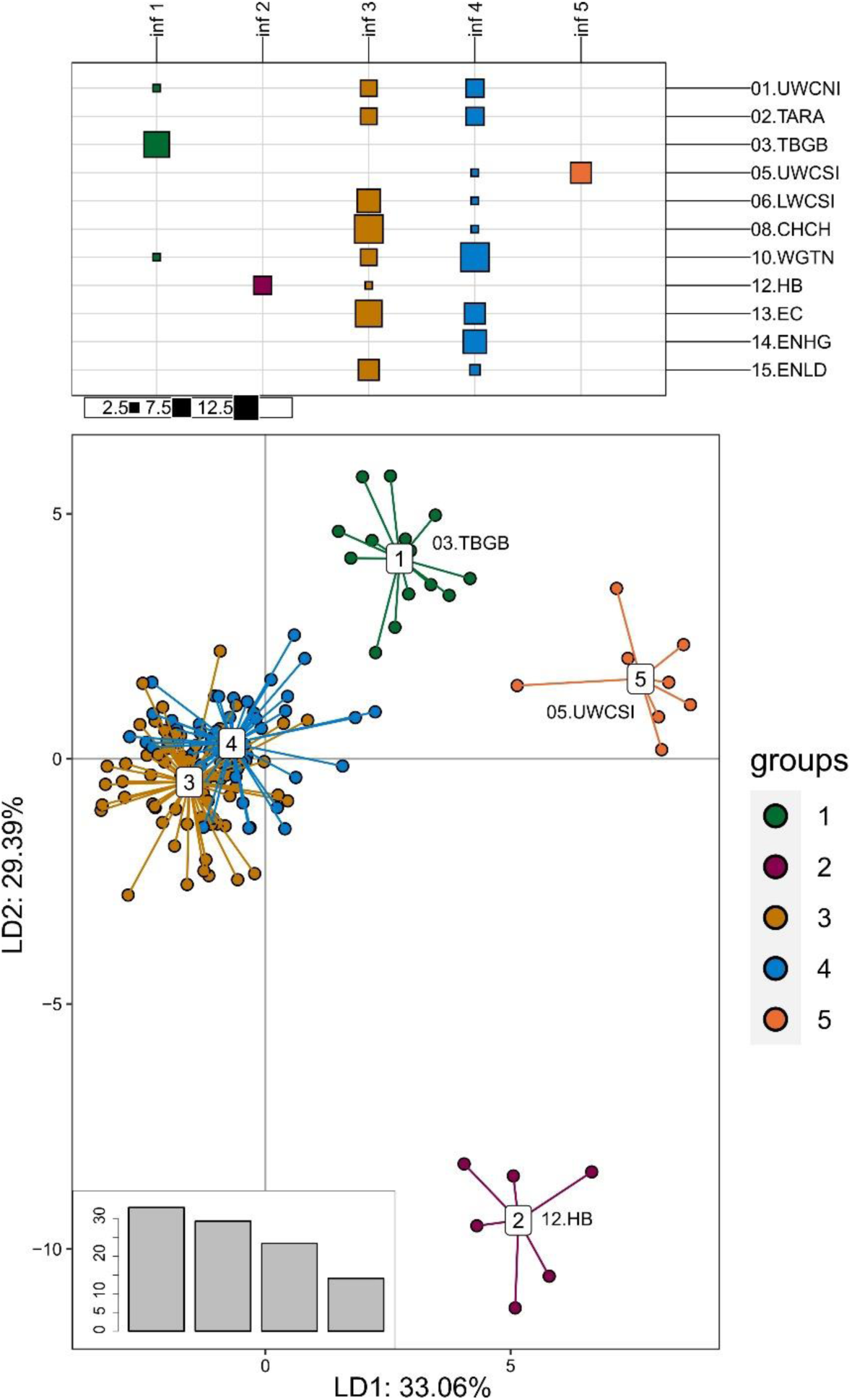
Discriminant analysis of principal components of the second adaptive SNP dataset that include 61 loci from 140 tarakihi. Top: results of the *K-*means clustering analysis, with colours corresponding to inferred groups (inf). Bottom: Projection of the DAPC based on inferred groups. Bottom left: Discriminant analysis eigenvalues. Sampling location codes as referred to in Table 1.

Pairwise weighted mean *F*_ST_ computed on the neutral dataset always showed high and significant values when comparing the Tasmania location with the New Zealand locations (*F*_ST_ = 0.0054– 0.0073, *P* ≤ 0.05 after false discovery rate correction), indicative of a clear genetic differentiation between tarakihi from Australia and New Zealand (Supplementary Figure 12). When comparing only populations from New Zealand (Figure 8), the *F*_ST_ values were globally low (*F*_ST_ = 0–0.0022), indicating a lack of overall sub-structure. However, some of the pairwise *F*_ST_ comparisons among locations were significant (*P* ≤ 0.05 after false discovery rate correction): in particular Wairarapa, which was significantly different to four other locations: Upper West Coast of South Island, Taranaki, East Northland, and East Cape (*F*_ST_ = 0.0012–0.0021). The only other significant differentiation was between Wellington and East Northland/Hauraki Gulf (*F*_ST_ = 0.0004). The dendrogram based on *F*_ST_ separated Wairarapa from all other New Zealand locations, which were split into two further clades separating Upper West Coast of South Island, Taranaki, and East Northland from the rest of the locations.

**Figure 8.**
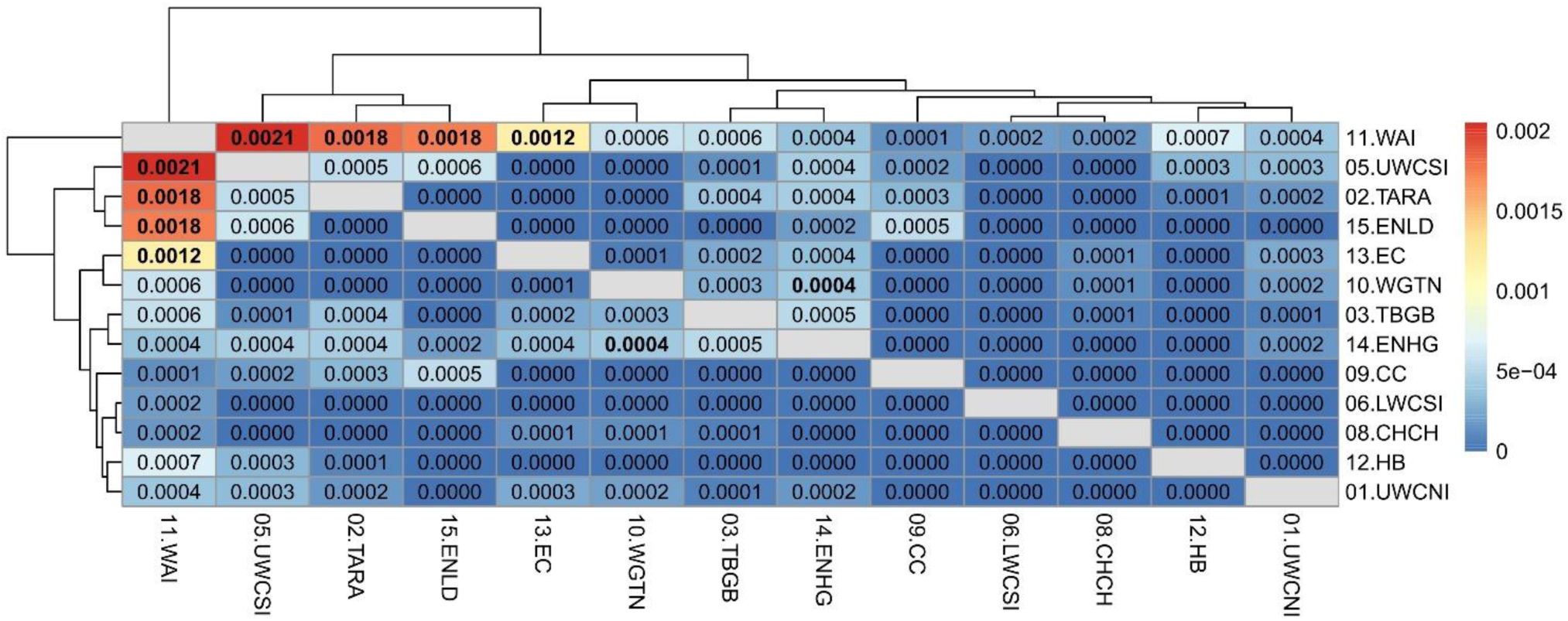
Heatmap of pairwise weighted *F*_ST_ estimates (corresponding to the values above and below the diagonal) among sample locations of the neutral SNP dataset, minus Tasmania. The dendrogram shows the inferred relationship between sample locations. Significant *p*-values (≤ 0.05 after false discovery rate correction) are in bold.

The relatively high divergence between Tasmania and New Zealand locations was also apparent in the adaptive dataset (Supplementary Figure 13) with all *F*_ST_ values being significant and ranging between 0.5232 and 0.5532. The mean pairwise *F*_ST_ values among New Zealand samples was also higher than in the neutral dataset (*F*_ST_ = 0.0251–0.1931) and all values were highly significant (*P* < 0.01 after false discovery rate correction) (Figure 9). This was expected since the dataset was composed of outlier SNPs with the highest *F*_ST_ values only. Wairarapa and Cape Campbell were the most divergent from the rest of the locations (*F*_ST_ = 0.1096–0.1931), followed by East Northland/Hauraki Gulf and Hawke’s Bay (*F*_ST_ = 0.0435–0.1832).

**Figure 9.**
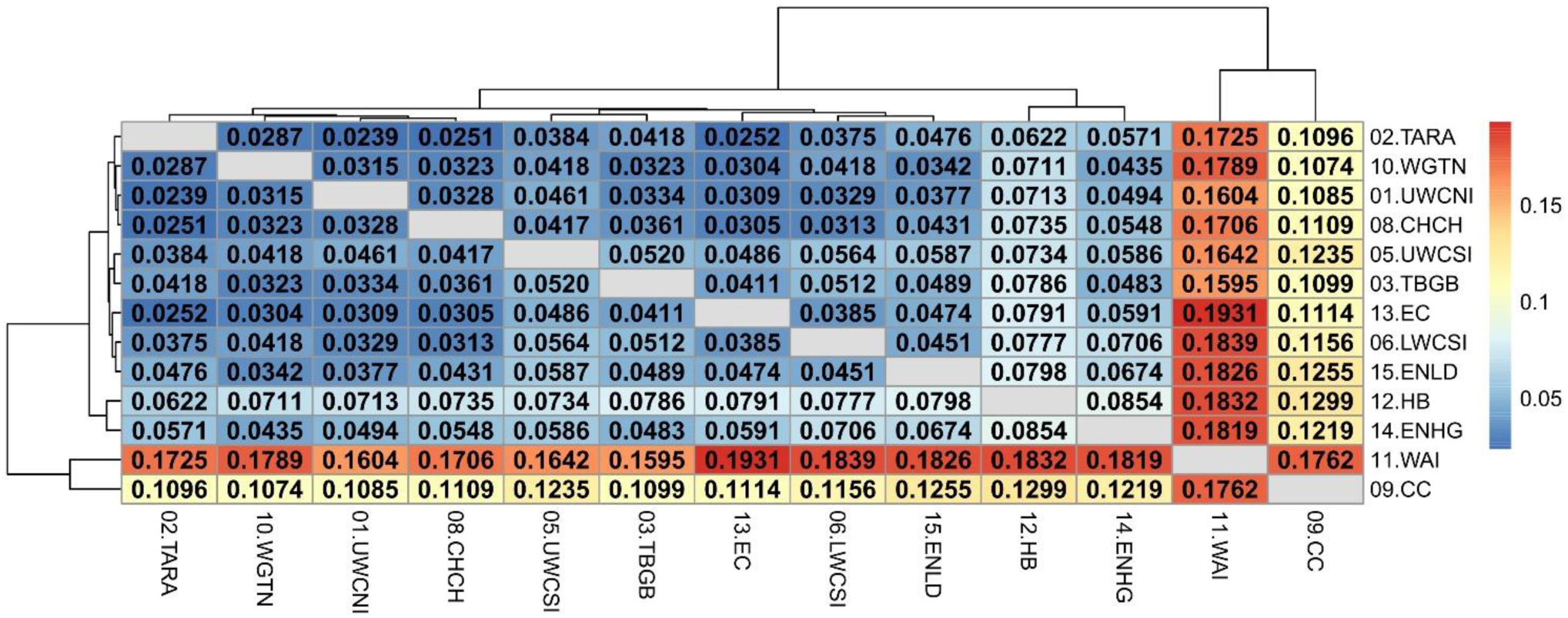
Heatmap of pairwise weighted *F*_ST_ estimates (corresponding to the values above and below the diagonal) among sample locations of the adaptive SNP dataset, minus Tasmania. The dendrogram shows the inferred relationship between sample locations. Significant *p*-values (< 0.01 after false discovery rate correction) are in bold.

FastSTRUCTURE did not detect any significant groupings in the neutral dataset, even including samples from Tasmania: the number of populations that best explained the structure was one. No significant pattern of isolation by distance was detected among New Zealand tarakihi locations when using a matrix of least-coast distance restricted to ocean travel on the neutral dataset (*R* = 0.260, *P* = 0.414).

### 3.5 Genotype-environment association analysis

Out of the eight longest scaffolds, only scaffold 1 was significant for its respective RDA model (*P* = 0.022). Moreover, for this model, only the first axis was significant (*P* = 0.01, with *P* > 0.1 for all other axes). Interpretations of the results were thus restricted to axis 1 of the RDA of scaffold 1. The adjusted *R*^2^ of the model was 0.05, and the variance inflation factors (vegan v2.5.7 function `vif.cca`) of the six predictor variables were all below 1.6, indicating that there was no multicollinearity among them. The projection of the SNPs, samples, and environmental variables on the first two axes shows that the sampling locations were effectively discriminated (Figure 10). Moreover, all samples projected in negative values of axis 1 were either from North Island or TBGB. Conversely, samples projected in positive values of axis 1 included all South Islands locations (except TBGB), all samples from Wairarapa and Chatham Islands, and a few samples from East Cape and Hawke’s Bay close to the center.

**Figure 10.**
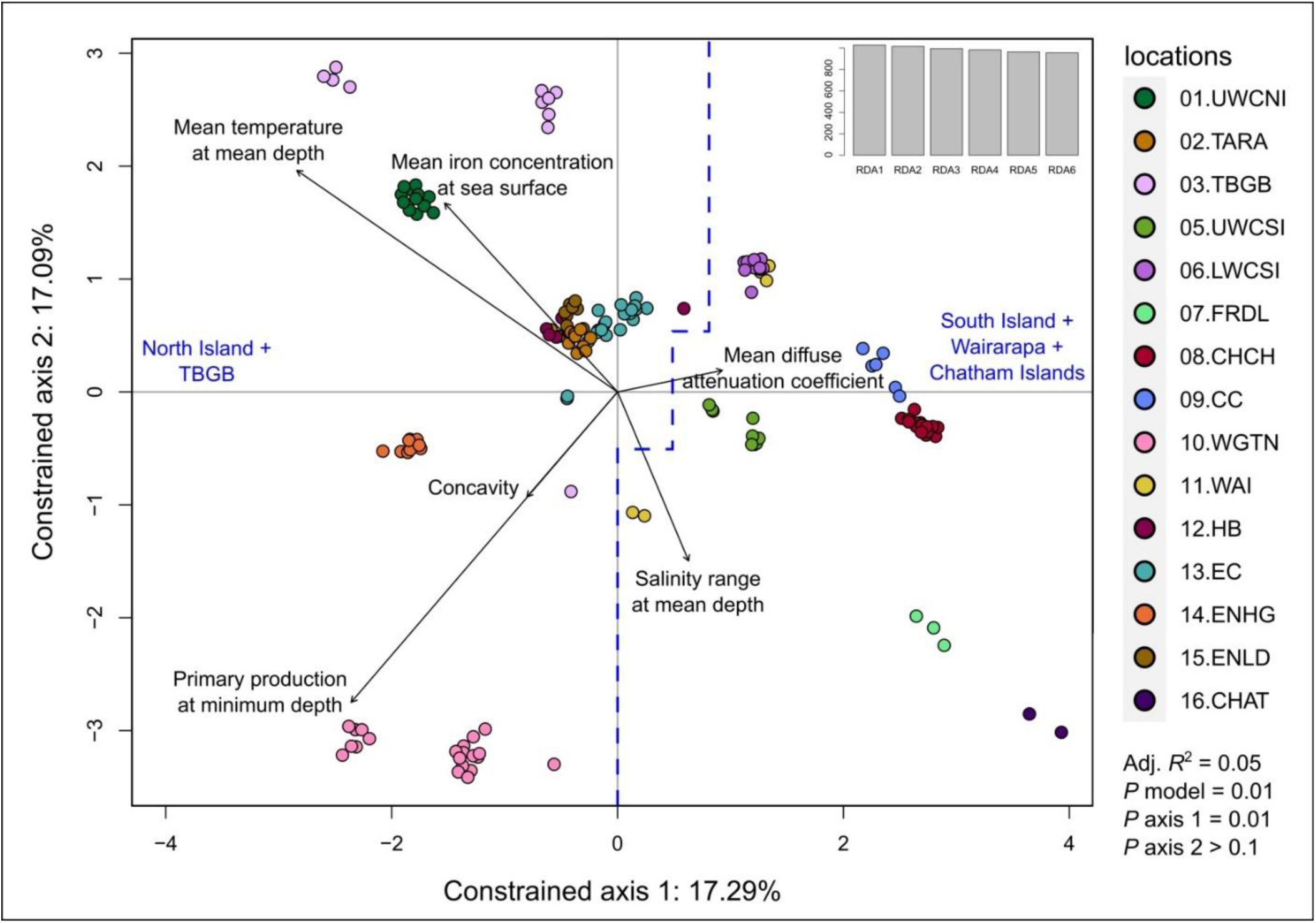
Redundancy analysis (RDA) performed on the quality-filtered SNP dataset from scaffold 1, restricted to adult tarakihi from New Zealand. This included 108,903 loci from 156 tarakihi. The two constrained axes show samples from 16 localities in relation to six lowly correlated environmental variables (black arrows). All samples on the left of the blue dashed line are from North Island and Tasman Bay/Golden Bay, while all samples on the right are from South Island, Wairarapa and Chatham Islands. Top right: Eigenvalues. Sampling location codes as referred to in Table 5.1.

Selection of outliers on the end tails of the loading distribution resulted in the identification of 55 candidate loci for local adaptation (Supplementary Table 3). Out of these 55 loci, 47 most strongly correlated with mean temperature at mean depth, five with mean primary production at minimum depth, two with mean iron concentration at the sea surface, and one with salinity range at mean depth. Thirty-two were located inside gene coding regions (with two pairs of loci being located in the same genes, *mindy3* and LOC111664994), eight were located in simple repeat regions, 14 in unannotated regions (most of them surrounded by highly-repetitive regions), and one in an unannotated, putatively transcripted region. A search of the Gene Ontology terms available for the same genes in zebrafish (*Danio rerio*) (Supplementary Table 4) showed that the most common associated GO terms were integral components of membranes (in four genes: transmembrane protein 67, glycoprotein endo-alpha-1,2-mannosidase-like protein, thrombospondin type-1 domain-containing protein 7A, and chondroitin sulfate proteoglycan 5a) and cilium assembly (in three genes: transmembrane protein 67, inositol polyphosphate-5-phosphatase B, and Zgc:171454 protein).

## 4 Discussion

This study investigated the neutral and adaptive stock structure on an expansive marine species that supports an important inshore commercial fishery. To detect environmental drivers associated with genetic structure, this study also applied gene-environment association analyses to gain insights into the selective forces acting on this species.

### 4.1 Genetic diversity

Tarakihi in New Zealand and Tasmania displayed similar levels of heterozygosity across their range (Supplementary Figure 3). The differences in heterozygosity observed in some individuals did not depend on the location but rather the sequencing depth, which means that a proportion of the genetic variation might not have been detected in the 14 lower coverage individuals. A plateau in heterozygosity seems to be reached when the mean coverage depth was > 7–8x, which indicates that the totality of the relevant genetic variation that could be captured at this level has been captured. The slightly higher mean heterozygosity for the neutral dataset could indicate that some of the filtered out outlier loci had a higher proportion of homozygotes, with some alleles possibly restricted to only a few populations in the dataset. A small fraction of less variable sites might also have been filtered out during the pruning step, due to deviations from Hardy-Weinberg and/or linkage equilibrium. The observed heterozygosity per individual is simply calculated as the proportion of heterozygous loci for that individual against the background of loci that are polymorphic in the dataset. It is difficult to say if this value is high in absolute terms. Comparison with results reported from other genome-wide SNP studies are not relevant, because contrary to e.g. mitochondrial markers, the observed loci are not necessarily directly comparable since the proportion of loci that are invariant across all individuals in each dataset is typically unknown (Gruber et al., 2021). A similar study using whole-genome resequencing in the Australasian snapper (*Chrysophrys auratus*) in New Zealand found a higher average heterozygosity of c. 0.20 for neutral loci (Oosting, 2021), while the snapper populations in New Zealand are smaller with supposedly lower connectivity among stocks than tarakihi.

Based on the same SNP dataset, it was clear that the heterozygosity in king tarakihi is lower than that of tarakihi (Ho = 0.06, Supplementary Figure 3B). This is expected from a population with a much smaller size and is consistent with recent findings of lower genetic diversity and smaller historical and current population size for this species (Papa et al., 2021a). However, since the reference genome was a tarakihi, it is possible that the heterozygosity values of the king tarakihi specimens were underestimated due to species-specific genome mapping success. This clear cut difference in heterozygosity was strong evidence that GBK had been misidentified when captured on the field, with a value corresponding to tarakihi and not king tarakihi (Ho = 0.12, Supplementary Figure 3B). The high number of fixed mutations between tarakihi and king tarakihi (84,144 fixed allelic differences) is additional strong evidence of the separate status of these two organisms (Smith et al., 1996, 2008; Burridge, 1999; Papa et al., 2021a). As a comparison, there are only three fixed allelic differences between tarakihi from Tasmania and New Zealand. These fixed mutational differences could be used as the basis for an SNP assay to differentiate tarakihi from king tarakihi in the field. The number of fixed SNPs among tarakihi in New Zealand locations was very low overall. Even for the Chatham Islands, where the number of mean fixed alleles with all New Zealand populations was 145, there were only five fixed alleles compared to the rest of New Zealand. This result could be due to undersampling because of the very low number of samples from Chatham Islands (*n* = 2).

### 4.2 Linkage disequilibrium

Linkage disequilibrium values were overall low in tarakihi, with mean *R*^2^ values never exceeding 0.2 (Figure 4). The decay rate was also very fast, with a plateau reached between 500 and 1,500 bp in all scaffolds. Fast decay of linkage disequilibrium is expected to be common for marine fishes, due to them maintaining very high effective population sizes over long periods of time (Hemmer-Hansen et al., 2014). Indeed, tarakihi effective population size has been historically very high, with a current estimation of about 100 million individuals (Papa et al., 2021a). However, linkage disequilibrium can also be impacted by admixture, mutation rate, founder effect, inbreeding and selection. Linkage disequilibrium values for genome-wide datasets are seldom available for marine organisms, but the results reported here are on par with values reported for European eel (*Anguilla anguilla*, complete LD below 10 kb) and Atlantic cod (*Gadus morhua*, complete LD decay below 10 centimorgans) by Hemmer-Hansen et al. (2014). This is also consistent with results reported from Australasian snapper, with a complete LD decay at 1,500 bp (Oosting, 2021).

Including the king tarakihi specimens in linkage disequilibrium analyses resulted in a slightly steeper disequilibrium decay and higher, more uniform *R*^2^ values along the scaffolds (Supplementary Figure 4). This could be indicative of an overall higher linkage disequilibrium in king tarakihi, due to a combination of small and recent stable population size. This could also indicate the presence of structural differences in the king tarakihi genome that would result in the physical distance between the bases being both underestimated and overestimated when aligned to the tarakihi genome, resulting in the detection of high linkage disequilibrium between relatively distant nucleotides (and inversely). Comparing results by aligning the king tarakihi SNPs to a contiguous king tarakihi genome assembly would be a good way to test for this.

### 4.3 Neutral genetic differentiation with king tarakihi and Australia

The strong genetic differentiation between tarakihi and king tarakihi is now well established (e.g. Supplementary Figure 3 and Figure 5). King tarakihi is thus a strong candidate for a formal taxonomic description as a separate species (tentatively *N. rex*) (Smith et al., 1996; Roberts et al., 2015, 2020; Papa et al., 2021a). However, king tarakihi is still currently reported and managed as part of TAR1.

Evidence for a partial lack of connectivity between Tasmanian and New Zealand tarakihi is very strongly supported by our results. A significant level of genetic divergence between these two areas (*P* ≤ 0.01) was detected by both AMOVA and pairwise *F*_ST_ analyses (Table 2, Supplementary Figure 12). Tasmanian individuals were always clustered together and separated from the New Zealand samples. These results support similar findings of trans-Tasman differentiation for tarakihi based on various genetic markers (Richardson, 1982; Elliott and Ward, 1994; Grewe et al., 1994), which microsatellites likely failed to detect (Burridge and Smolenski, 2003). It can be reasonably extrapolated that this genetic divergence to New Zealand would also be found in samples analysed from other locations in the Australian range, meaning there are likely two separate reproductive stocks, one in Australia and one in New Zealand. It would be useful to compare samples taken from additional locations in Australia: some genetic studies have reported a genetic distinction between fish stocks from west and east Australia even when the gene flow between Australia and New Zealand was putatively high. This is the case for e.g. gemfish (*Rexea solandri*) (Colgan and Paxton, 1997), kingfish (*Seriola lalandi*), (Nugroho et al., 2001; Miller et al., 2011), and mako shark (*Isurus oxyrinchus*) (Corrigan et al., 2018).

### 4.4 Neutral genetic structure in New Zealand tarakihi

The analysis of 166,022 neutral genome-wide SNPs conducted in this study indicates that tarakihi have a panmictic genetic population structure throughout their distribution around mainland New Zealand. This may also include the Chatham Islands, although the number of samples for this location is too low to test this. No obvious genetic structure related to sampling locations or management areas were detected by any of the methods used (AMOVA, PCA, DAPC, and FastSTRUCTURE) and no significant isolation by distance was detected either. The only significant genetic differences detected were through the pairwise *F*_ST_ between Wairarapa (south-east of North Island) and four other locations: Upper West Coast South Island, Taranaki, East Northland, and East Cape (Figure 8).

This result is difficult to interpret in terms of geographic stock structure since these four locations are situated at several distant areas around New Zealand and are separated from the Wairarapa by other non-significantly divergent locations (e.g. TBGB, Hawke’s Bay and Hauraki Gulf) (Figure 2). The divergence with East Cape and East Northland could be indicative of a complex, fine-scale migration pattern northward along the east coast of North Island, which would be partially concordant with the results from the mitochondrial study (Papa et al., 2021a). The divergence of Wairarapa compared to Taranaki and Upper West Coast of South Island could be due to a partial lack of connectivity between Wairarapa and the west coast through Cook Strait. However, this seems unlikely since no significant genetic divergence was detected between Wairarapa and Upper West Coast of North Island or Lower West Coast of South Island. It is worth noting that the genetic divergence found between Wairarapa and these four locations, while still significant after fdr correction, is very low (*F*_ST_ = 0.0012–0.0021). Moreover, while all Wairarapa samples have been sequenced at a good depth and have a “normal” (i.e. expected) level of heterozygosity (Supplementary Figure 3), and therefore the relevant individual genetic variation can be reasonably thought to have been captured, the number of individuals from this location is the lowest included in the neutral dataset (*n* = 5 individuals only). The detected genetic divergence might thus be an artefact due to the low number of samples for this location, causing individuals to have a high proportion of fixed allelic differences because the totality of the allelic variation in that location has not been sampled. However, simulation studies have shown that genetic differentiation measured by *F*_ST_ can still be accurately estimated in populations as small as *n* = 4–6 when the number of SNPs is above thousands (Willing et al., 2012). No comparison with the study from Gauldie & Johnston (1980) could be done since they did not include samples from around Wairarapa.

This overall lack of population genetic structure for tarakihi around New Zealand is concordant with results from Papa et al. (2021a) where no evidence of genetic structure was detected for the overall New Zealand area, including Chatham Islands. These results appear to consistently suggest that there is one panmictic tarakihi stock in New Zealand, with very high spatial connectivity and no spatial reproductive isolation. This finding is concordant with the characteristics of a species with a high capacity for dispersal, reproduction output, and effective population size, and would also explain the failure to detect any genetic structure among Australian stocks either, despite numerous attempts (Richardson, 1982; Elliott and Ward, 1994; Grewe et al., 1994; Burridge and Smolenski, 2003).

### 4.5 Adaptive genetic structure

The acquisition of a first and then a second set of putatively adaptive outlier SNPs resulted in the detection of fine-scale genetic structure that was not observed in the neutral tarakihi genetic dataset (Figures 6&7, Figure 9). Interestingly, the presumably adaptive genetic variation detected always discriminated groups at the sampling location level, rather than clustering groups at a broader geographic level (e.g. management area, coast, island). The first set of 389 presumably adaptive SNPs discriminated Tasmania from New Zealand, meaning that the genetic divergence observed between these two stocks is likely due to both physical isolation and local adaptation. They also discriminated Wairarapa and Cape Campbell from all other New Zealand locations (Figure 6). While Wairarapa and Cape Campbell are geographically close (south-east of North Island and north-east of South Island, Figure 2) it is not obvious if they should be considered together as one or two different adaptive stocks separated from New Zealand: although they formed two different clusters on the PCA and DAPC (Figure 6 & Supplementary Figure 10), most of the variation of both groups were on the axis 2 of the DAPC (Figure 6) and on both the axes 2 and 3 of the PCA (Supplementary Figure 10). Moreover, the dendrogram inferred on the pairwise *F*_ST_ clustered the locations together as outgroup of the rest of New Zealand, even though the *F*_ST_ value between them (*F*_ST_ = 0.1762) was one of the highest among all pairwise locations (Figure 9).

Excluding Tasmania, Wairarapa and Cape Campbell improved the statistical power of the outlier method. With this second dataset of 61 putative adaptive SNPs, the *K*-mean clustering analysis was able to also discriminate Upper West Coast South Island, Hawke’s Bay, and TBGB without any *a priori* grouping. Interestingly, although TBGB and Upper West Coast South Island are both on the west coast of South Island, these three putatively adaptive stocks seem to be actually separated from each other and from the rest of New Zealand (Figure 7).

Wairarapa, Cape Campbell, Hawke’s Bay, Upper West Coast South Island, and TBGB were found to be genetically divergent from each other and from all of the remaining New Zealand locations, which includes sites as far apart as East Northland, Christchurch, and lower West Coast South Island. If these outlier-based stocks are indeed adaptive, this would indicate that there is no broad separation of adaptive genetic stocks around New Zealand, but rather that the tarakihi population displays some very fine-scale local adaptation to specific areas around New Zealand, that are not, at first look, directly linked to e.g. temperature or depth. This adaptive genetic variation could be school-specific, i.e. representative of genetically adapted small groups, rather than due to large-scale environmental factors. Interestingly, Gauldie & Johnston (1980) also found genetic differentiation between lower and upper west coast of South Island, although the sampling locations were a bit further apart (Jacksons Bay and Westport). While they also detected a genetic break between TBGB and Cape Campbell, the former was thought to be part of a broader western genetic stock including Upper West Coast South Island and Taranaki and the latter an eastern one including Christchurch and East Cape. Given these results and the small sample size of some of these locations (especially Wairarapa, *n* = 5 and Hawke’s Bay, *n* = 7), it is legitimate to question if these observations are statistical artefacts. The outlier method used here is particularly suited to detect loci under heterogeneous selection and local adaptation (Whitlock and Lotterhos, 2015). Although the two SNP datasets obtained with this method are only putatively adaptive, they are very strong candidates for adaptation because the parameters used to detect them were quite stringent: only loci with *q* < 0.05 and He > 0.1 were retained, the first to greatly minimize the risk of false positives and the second to discard low-frequency alleles that do not fit the neutral *F*_ST_ distribution used to find outliers. Moreover, the OutFLANK method does not rely on an *a priori* population model, and is very robust against false positives: in fact, when the number of individuals per location is low, OutFLANK tends to lose power and will produce more false negatives instead of false positives (Whitlock and Lotterhos, 2015). Thus, it is likely that the adaptive divergence observed is biologically relevant and not an artefact due to the small sample size of some locations.

### 4.6 Temperature-associated selective cline

Contrary to the PCA and DAPC performed on the neutral and outlier-based adaptive SNP datasets, the RDA of the reduced quality-filtered SNP dataset was very effective at discriminating samples based on localities (Figure 10). Given that only the first axis of the RDA on scaffold 1 was significant, and that 47 out of 55 (85%) of the candidate environmentally-adapted loci were most strongly correlated with mean temperature at mean depth, the variation observed on axis 1 appears to be driven by adaptation to temperature, or to any other environmental variable strongly correlated with temperature (e.g. salinity, pH, dissolved oxygen concentration). Moreover, the samples were ordinated following a latitudinal gradient, which is directly related to temperature on the continental shelf, where tarakihi occur (Supplementary Figure 14). Only TBGB and Wairarapa are not projected with, respectively, the South and the North Island (Figure 10). This is explained by the fact that TBGB is actually situated further north than Wairarapa. This means that the observed genetic variation is directly related to temperature (and thus latitude) rather than a theoretical differentiation between the North and the South Island driven by e.g. a dispersal barrier in Cook Strait. This was verified with a linear regression analysis that showed that there was a significant correlation between the ordination on the RDA axis 1 and both the mean temperature and the latitude at sampling locations (Pearson *R*^2^ = - 0.65, *P* < 0.001 for both variables, Supplementary Figure 15).

### 4.7 Fisheries management implications

It is now well established that the Quota Management Area boundaries for tarakihi in New Zealand do not match the biological stock boundaries (Langley, 2018; Fisheries New Zealand, 2021). The Chatham Islands are usually considered a separate stock from the North and South Island because these two areas are geographically separated by deep water which is not usually inhabited by adult tarakihi (Morrison et al., 2014; Fisheries New Zealand, 2021). Although the main islands are split into several management areas (Figure 2), they are not thought to be accurate reflections of the reproductive stock boundaries. Both North and South Islands have been generally considered as one single stock due to the high capacity of post-larvae and adults for dispersal and the lack of evidence of genetic isolation (Morrison et al., 2014; Fisheries New Zealand, 2021). Some specimens tagged in the Kaikoura area on the east coast of South Island have been recaptured on the west coast of North Island (Hanchet and Field, 2001), suggesting that there is at least some level of connectivity between the west and east coasts. Recent studies on trends in age and size structure in TAR1, TAR2 and TAR3 found evidence that the east coast of North and South Island could be one continuous stock (McKenzie et al., 2017; Langley, 2018). The Canterbury Bight/Pegasus Bay area (east of South Island) seems to be the main nursery area for juveniles of the entire eastern stock. At the onset of maturity, some of the adult fish migrate northward along the east coast to East Cape, then up to the east Northland area. This scenario is backed up by the higher proportion of juveniles in Canterbury Bight/Pegasus Bay and the observed increase in the proportion of older fish from TAR 2, Bay of Plenty and east Northland (Langley, 2018). Subsequent studies have found a similar pattern for the west coast of New Zealand (Fisheries New Zealand, 2021). Observations about the relative strength of individual year classes and growth rates of older fish seem to indicate that Tasman Bay/Golden Bay (TBGB) could act as a nursery area for the entire west coast, which would constitute a single separated stock from the east coast, with a possible lack of connectivity between the upper west coast of North Island and the East Northland fisheries (Fisheries New Zealand, 2021).

There is thus strong preliminary evidence that tarakihi are constituted of two main stocks in New Zealand, however, the present analysis of neutral genomic variation does not support this hypothesis, since no genetic divergence was detected between the west and the east coast, for both islands, even when grouping them using AMOVA. Moreover, the weak genetic divergence that was detected between the west and east coast of the South Island using mitochondrial data (Papa et al., 2021a) could not be corroborated in this study. A similar situation occurred with the Australian stocks: the observation of no genetic structure in Australia reported by Burridge & Smolenski (2003) was in contrast with two studies that detected three stocks within the south-east of Australia based on otolith microchemistry (Thresher et al., 1994) and larval advection (Bruce, 2001).

When a test for genetic differentiation rejects the model of panmixia, it can usually be used with confidence as a biologically meaningful finding. However, it is more difficult to firmly conclude that stocks are demographically coupled whenever the test fails to reject panmixia (Waples et al., 2008). The demographic structure described by Langley (2018) might not be a reasonable conclusion from the present population genomic analyses because there may be a high enough level of connectivity between stocks to sufficiently homogenise genetic variation, but not enough to provide demographic coupling between areas. Some level of migration between the west and east coasts is highly likely, based e.g. on tagging studies (Hanchet and Field, 2001). Only a few migrants per generation are needed to homogenise most genetic loci, and a large effective population size reduces the speed at which differentiation occurs through the action of genetic drift. Furthermore, any demographic separation between stocks might have arisen too recently for a parallel pattern to also be seen in a test for genetic differentiation. The population genomics findings do not necessarily contradict the findings reported from age and size data. There is still the possibility that there is a western and an eastern stock that may only be partially reproductively isolated, and that some small level of gene flow still occurs and is enough to homogenise genetic variation.

The outlier-based, presumably adaptive differentiation reported here is unlikely to be directly useful for the delineation of fisheries management stocks at this stage, since the genetic variation appears to be highly localized and not reflecting any known broader stock boundaries (Figure 2). However, the genotype-environment association analysis hints at the possibility of a North-South adaptive cline in the stock related to water temperature (Figure 10, Supplementary Figure 15). The pattern of genetic variation found at these loci should be monitored through time to test whether changes in water temperatures have influenced the distribution of alleles over time. The tarakihi population may show an adaptive response to future climate and sea temperature changes, but the consequences of this for the resilience of the fishery is largely unknown.

While the level of population genomic resolution found in this study was high, the number of individuals sampled is relatively low, especially in some locations because very few samples could be obtained. Simulation studies have shown that genetic structure can sometimes be detected by only sampling a few individuals per population (c. 8–10) as long as the genome-wide SNP density is high (Jeffries et al., 2016; Nazareno et al., 2017). However, the power of clustering methods (i.e. K-means clustering for DAPC) can drop sharply when a small number of individuals is used (Waples and Gaggiotti, 2006), and given the high effective population size and the high level of gene flow in New Zealand tarakihi, a much larger DNA sampling and sequencing campaign might be necessary to provide fisheries managers with a definitive answer of the presence or absence of genetic stock structure among tarakihi in New Zealand.

## 5 Conclusion

This study is the first population genomics analysis of tarakihi (*Nemadactylus macropterus*) and one of the very first genome-wide analyses of a New Zealand marine species. The acquisition and subsequent filtering of a large SNP dataset allowed for the detection of a low but highly significant genetic differentiation between Tasmania (and thus putatively the whole of Australia) and New Zealand. No neutral genomic structure was detected among New Zealand locations, which means the genomic data did not support the hypothesis of two separate reproductive stocks on the west and east coast of New Zealand. A latitudinal adaptive cline strongly correlated to water temperature was found. The associated loci are strong candidates for further investigation to identify potential functional adaptive role.

Tarakihi is a commercially important fishery but it has been reported as declining. Implementation of routine genomic sampling could enhance spatial and temporal genomic resolution. This could be implemented using a long-term, standardised, well-archived genetic-based tagging program (Mace et al., 2020; Papa et al., 2021b). Integrating results obtained from genetic-tagging data with life history information and results obtained from other fisheries assessment methods will prove extremely valuable for the long-term sustainable use of this important fishery species.

## 7 CRediT authorship contribution statement

**YP:** Conceptualization, Methodology, Software, Validation, Formal analysis, Investigation, Resources, Data Curation, Writing - Original Draft, Writing - Review & Editing, Visualization. **MM:** Resources, Writing - Review & Editing, Supervision, Funding acquisition. **MW:** Writing - Review & Editing, Supervision. **PR:** Conceptualization, Resources, Writing - Review & Editing, Supervision, Project administration, Funding acquisition.

## 9 Acknowledgments

We are grateful to the following people and companies who contributed to this study. Collection of the tarakihi and king tarakihi specimens from phase 1 (2017–2018) was supervised by Cameron Walsh (Stock Monitoring Services Limited). Specimens were collected by fishing companies (Gisborne Fisheries, Star Fish Supply, Egmont Seafoods, Moana New Zealand, Hawke’s Bay Seafoods, United Fisheries, Talley’s Seafood, and Wellington Trawling) and Peter Young (Cruise Fiordland). Alex Halliwell (Victoria University of Wellington) provided assistance for the tissue collections and DNA extractions. The majority of samples from phase 2 (2019–2020) were collected by NIWA staff including Jeremy McKenzie, Dan MacGibbon, Helena Armiger, Jade Arnold, and Caoimhghin Ó Maolagáin. Samples from Australia were collected by Anne-Marie Hegarty and Matt Taylor (New South Wales Department of Primary Industries). Additional tarakihi tissue samples were collected by Tom Oosting (Victoria University of Wellington) at Gisborne Tatapouri Sports Fishing Club and Hawke’s Bay Sports Fishing Club fishing competitions. We are grateful to Alison Wilson for providing editorial feedback.

## 11 Data Availability Statement

The datasets generated for this study can be found in the Genomics Aotearoa repository [https://data.agdr.org.nz]. All bash and R scripts used for this study are available on GitHub on the following repository: https://github.com/yvanpapa/tarakihi_population_genomics.

