## Supplementary Material for "Genomic stock structure of the marine teleost tarakihi (*Nemadactylus macropterus*) provides evidence of fine-scale adaptation and a temperature-associated cline amid panmixia"

### 1 Supplementary Data

#### 1.1 Supplementary Figures

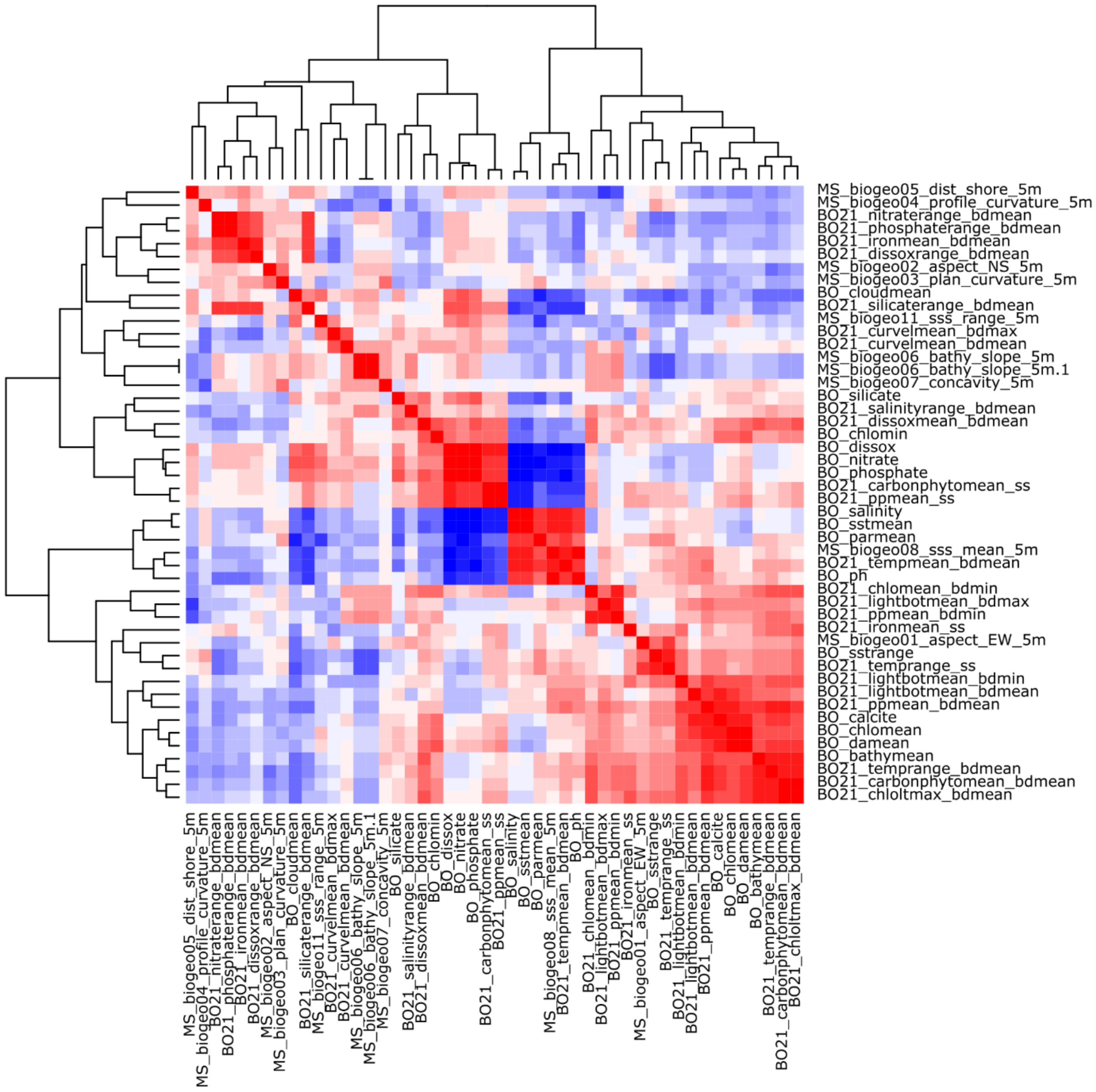


**Supplementary Figure 1.** Heatmap of Person correlation coefficients among the 48 environmental variables compiled in the second dataset.

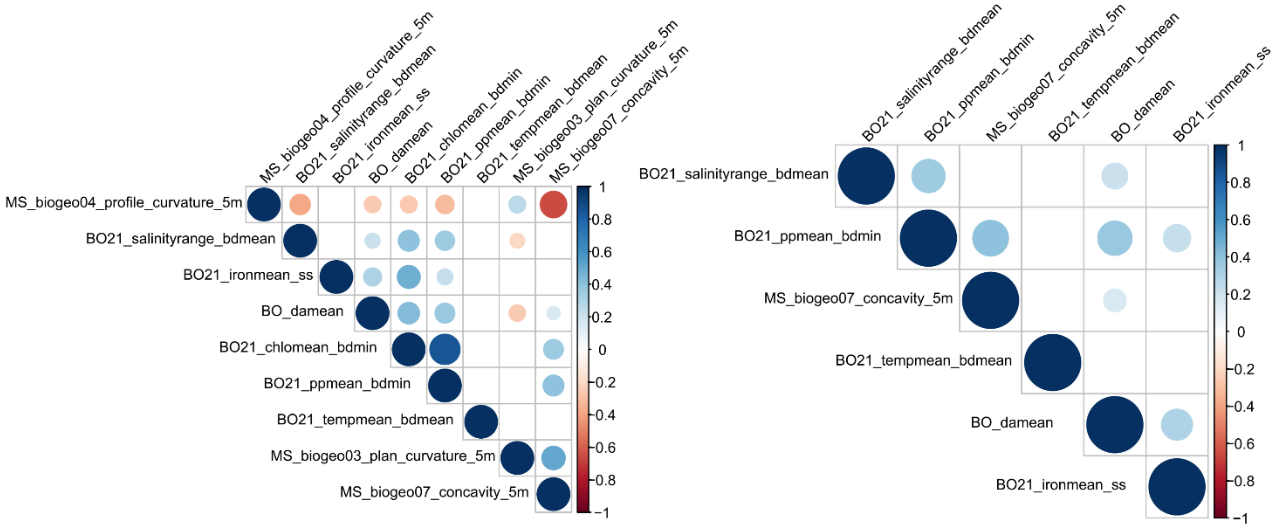


**Supplementary Figure 2.** Correlation plot of the environmental variables after discarding for correlation with mean temperature at mean depth (left), and the final dataset of six lowly correlated variables used in the locus-environment association analysis (right). Color scale and size of dots correspond to Pearson coefficient value. Dots are plotted only for significant values ($P \leq 0.05$).

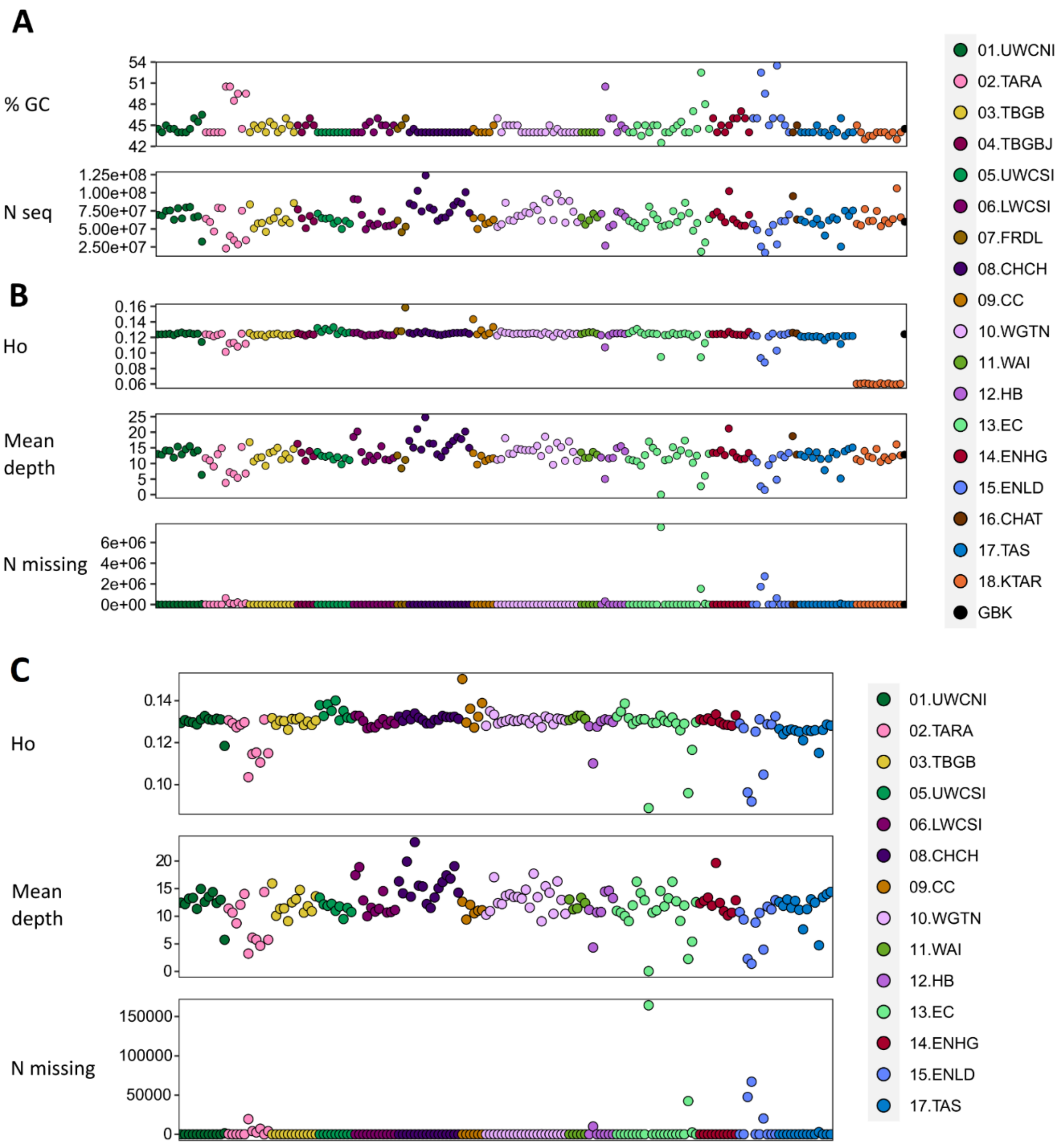


**Supplementary Figure 3.** Summary statistics computed on (A) trimmed Illumina reads, (B) quality-filtered SNP dataset, and (C) neutral SNP dataset. N seq: number of Illumina reads. Ho: mean observed Heterozygosity. N missing: number of missing sites. Each circle is an individual on the X-axis.

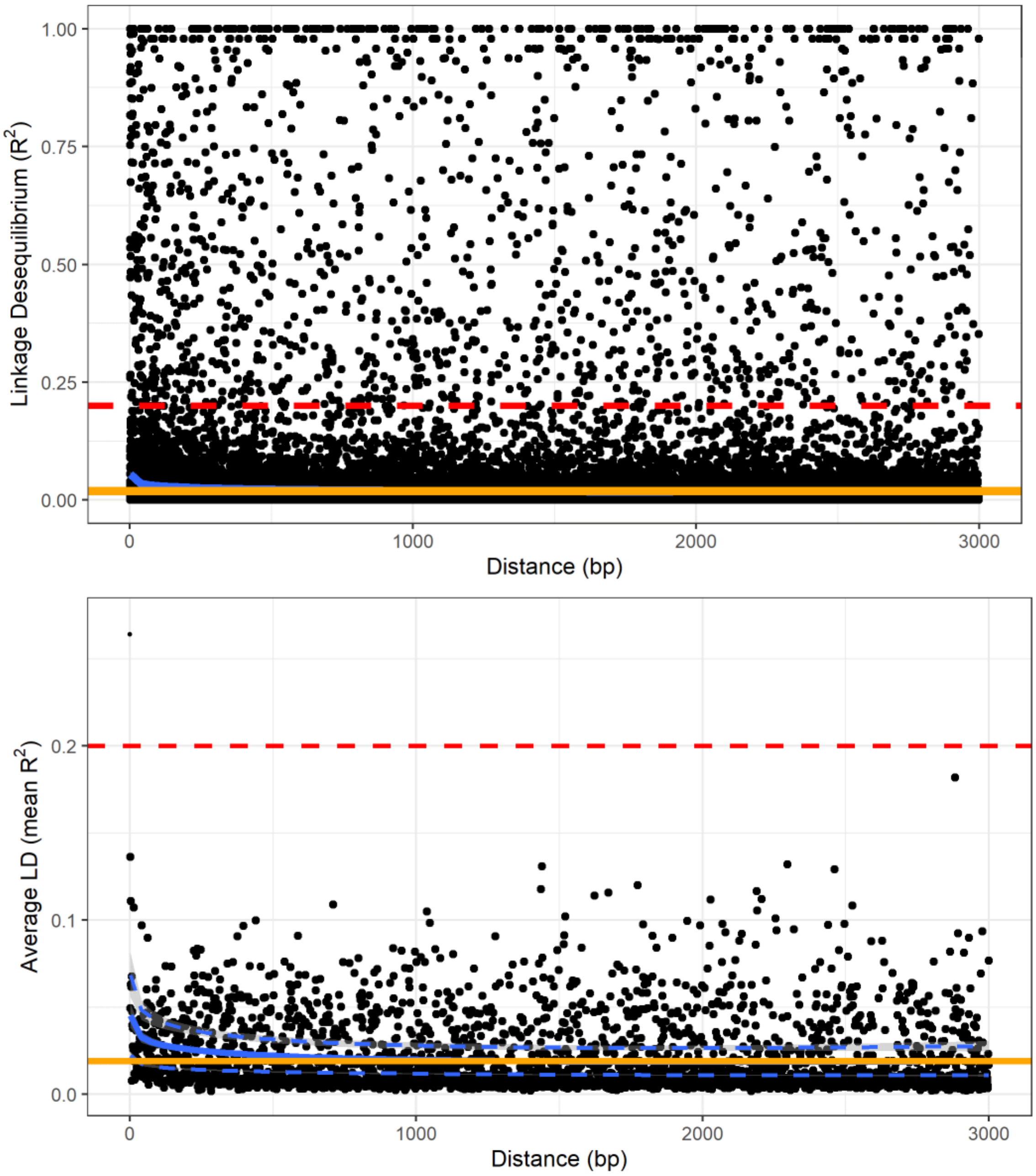


**Supplementary Figure 4.** Linkage disequilibrium decay over genetic distance on scaffold 1, calculated on the quality-filtered SNP dataset, including king tarakihi specimens. The horizontal red dashed line shows the threshold of 0.2 that is commonly applied to identify independent degradation of nucleotide sites. The orange line is the background level of linkage disequilibrium (intercept). The blue line is the trend of linkage disequilibrium decay fitted to the plot (minimum and maximum variance in dashed lines for the mean).

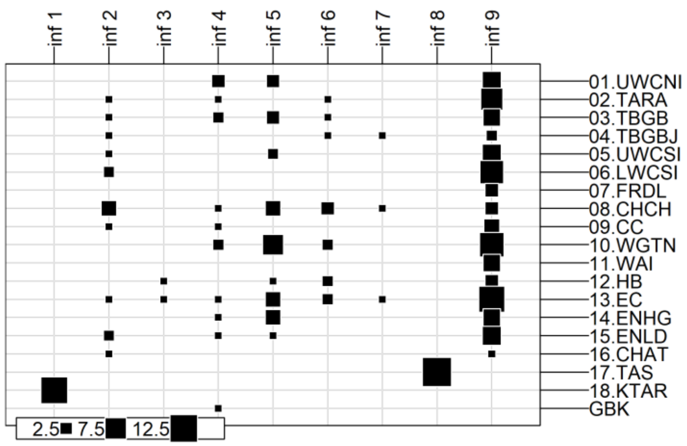


**Supplementary Figure 5.** Results of the *K*-means clustering analysis performed on the pruned SNP dataset that includes 183,443 loci from 188 tarakihi and king tarakihi.

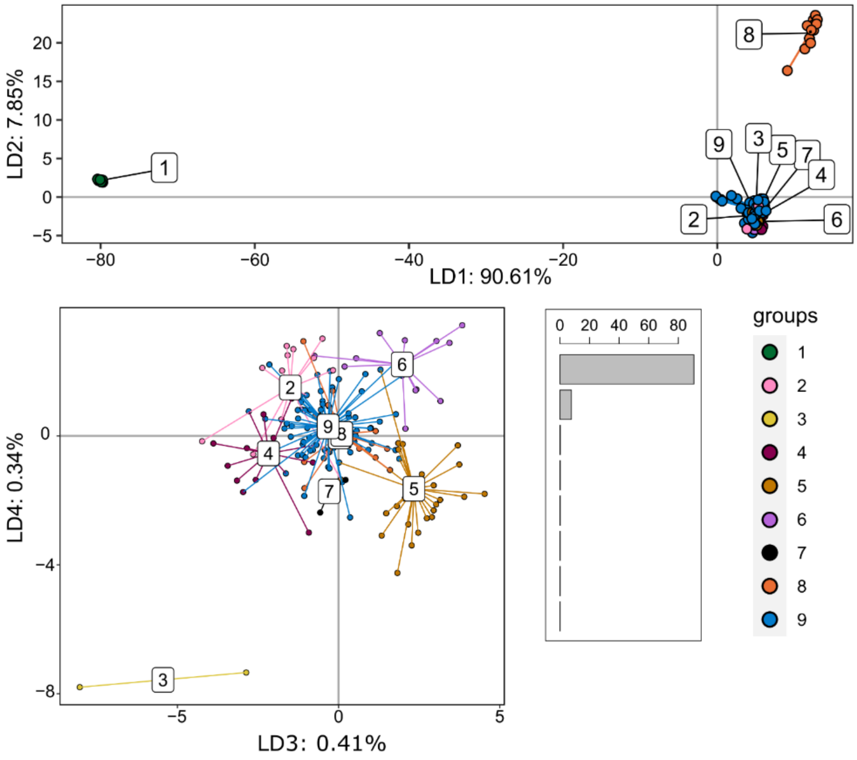


**Supplementary Figure 6.** Discriminant analysis of principal components of the pruned SNP dataset that includes 183,443 loci from 188 tarakihi and king tarakihi. Top: Axes 1 and 2. Bottom left: Axes 3 and 4. Bottom right: Eigenvalues. Groups correspond to the inferred groups in Supplementary Figure 4.

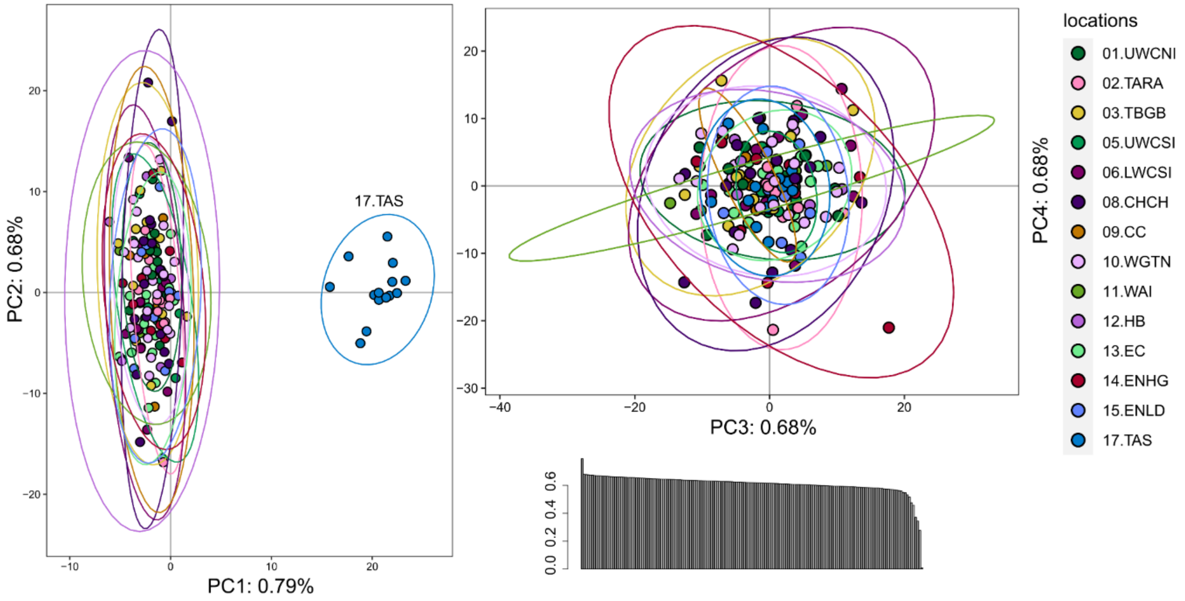


**Supplementary Figure 7.** Principal component analysis of the neutral SNP dataset that includes 166,022 loci from 165 tarakihi. Ellipses represent the 95% confidence intervals. Left: Axes 1 and 2. Top right: Axes 3 and 4. Bottom right: Eigenvalues. Sampling location codes as referred to in Table 1.

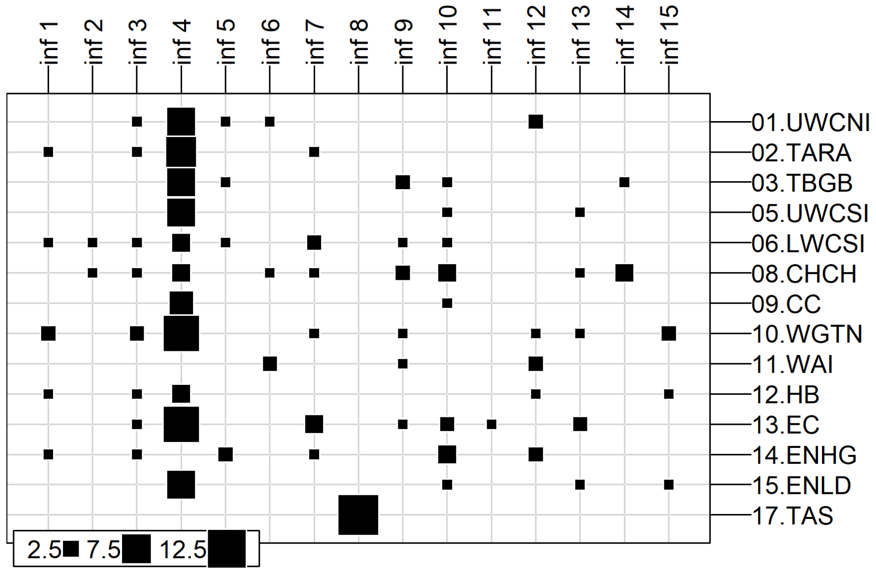


**Supplementary Figure 8.** Results of the *K*-means clustering analysis performed on the neutral SNP dataset that includes 166,022 loci from 165 tarakihi.

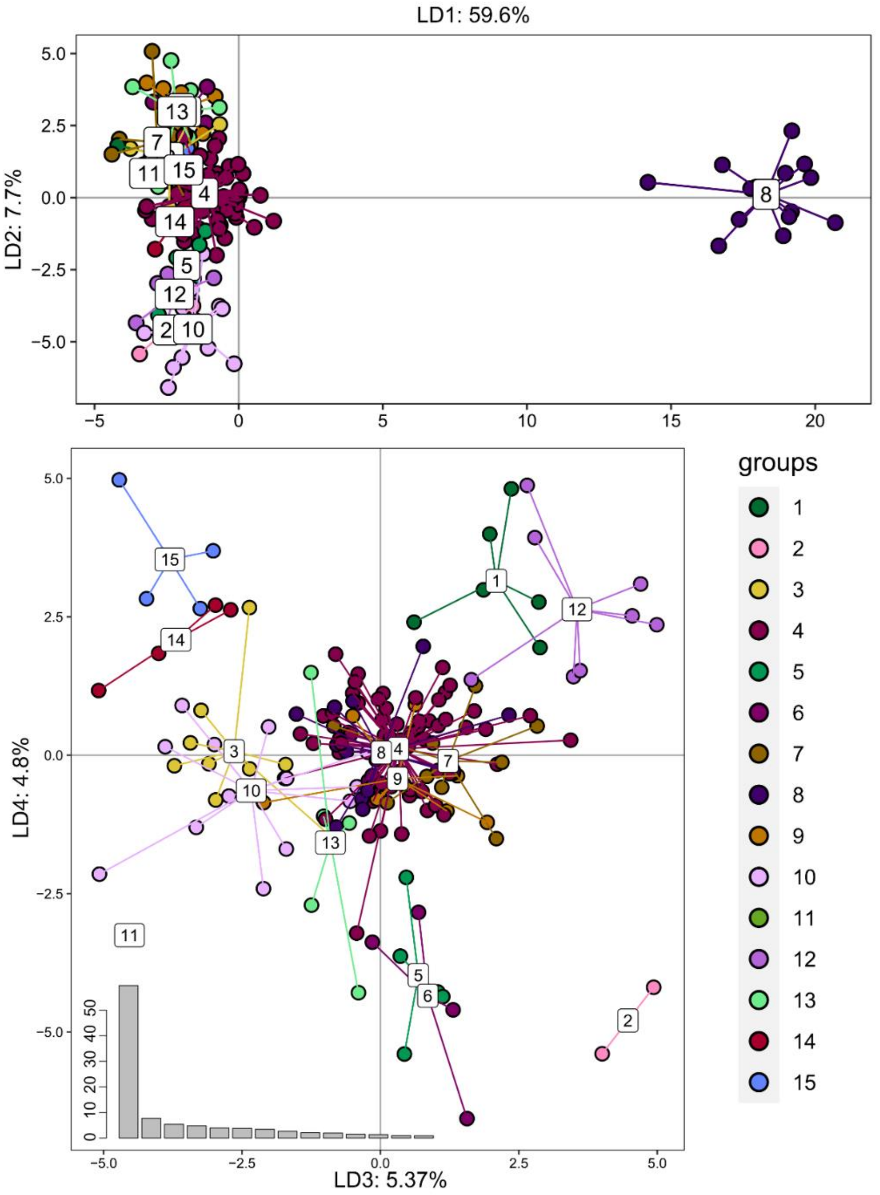


**Supplementary Figure 9.** Discriminant analysis of principal components of the neutral SNP dataset that includes 166,022 loci from 165 tarakihi. Top: Axes 1 and 2. Bottom left: Axes 3 and 4. Bottom left: Eigenvalues. Groups correspond to the inferred groups in Supplementary Figure 7.

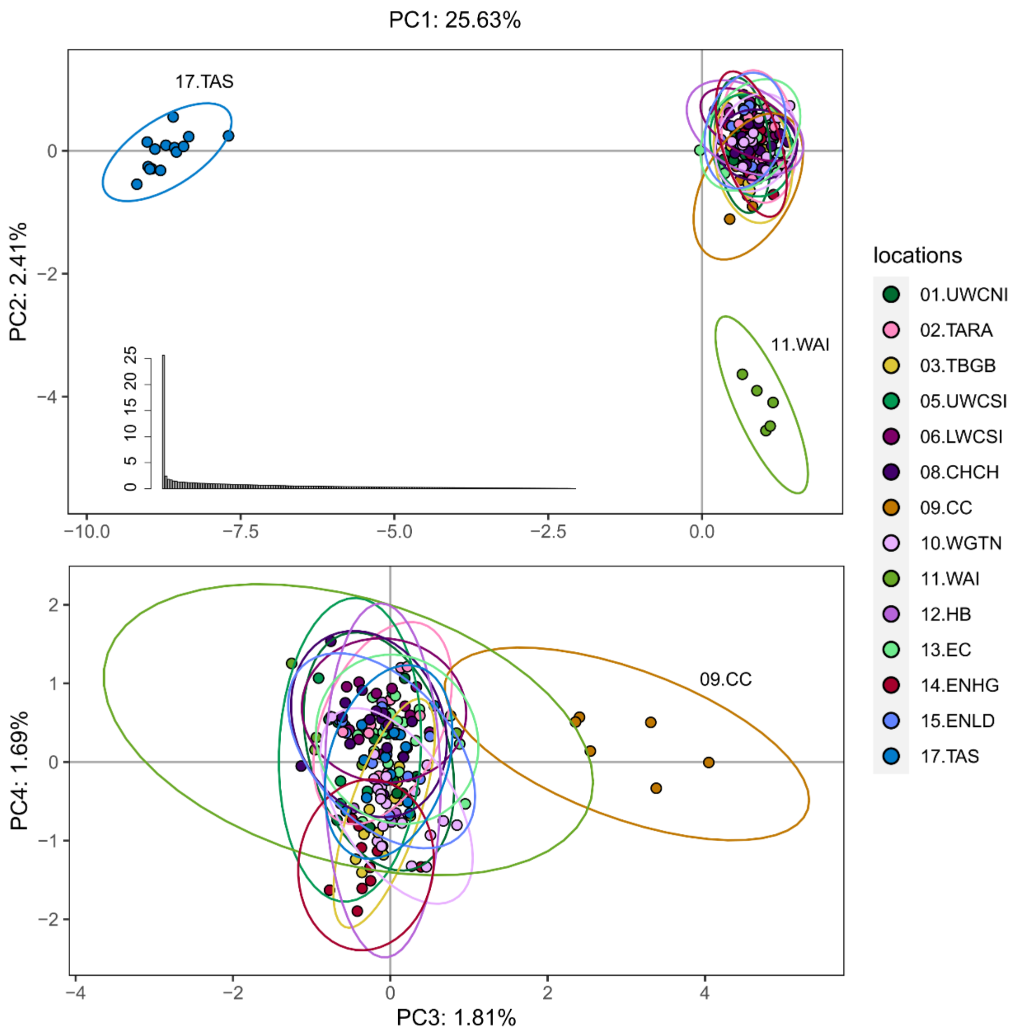


**Supplementary Figure 10.** Principal component analysis of the adaptive SNP dataset that includes 389 loci from 165 tarakihi. Ellipses represent the 95% confidence intervals. Top: Axes 1 and 2. Bottom: Axes 3 and 4. Middle: Eigenvalues. Sampling location codes as referred to in Table 1.

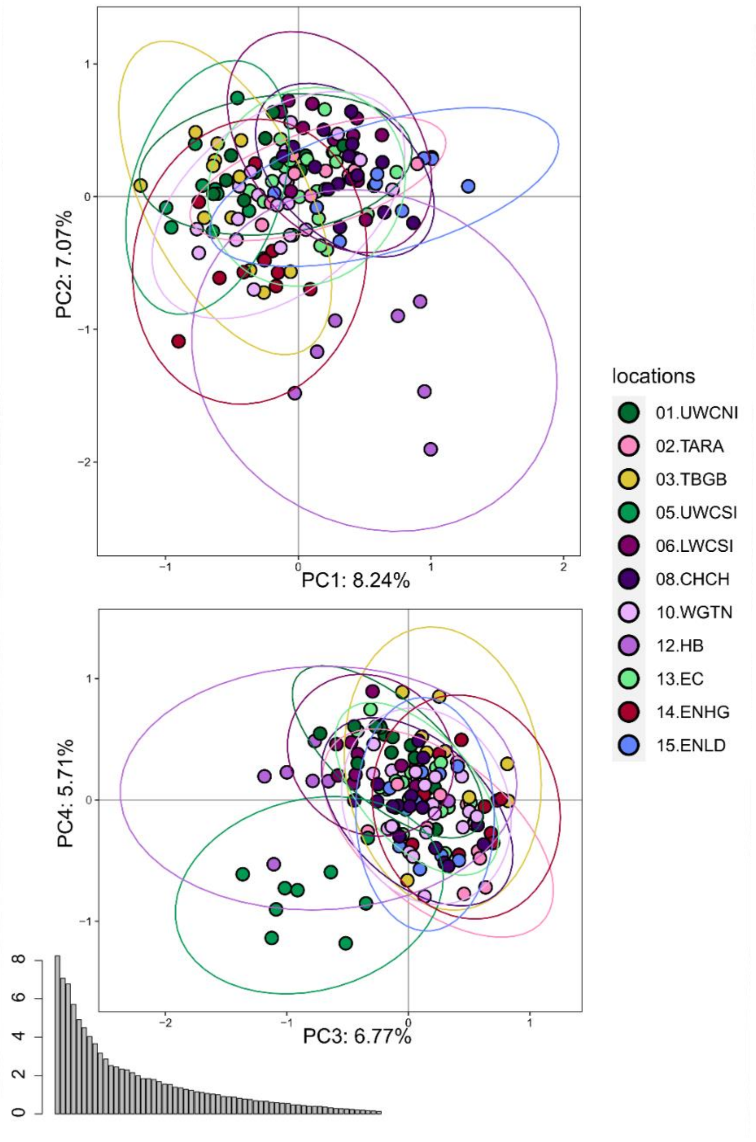


**Supplementary Figure 11.** Principal component analysis of the second adaptive SNP dataset that includes 61 loci from 140 tarakihi. Ellipses represent the 95% confidence intervals. Top: Axes 1 and 2. Bottom: Axes 3 and 4. Bottom left: Eigenvalues. Sampling location codes as referred to in Table 1.

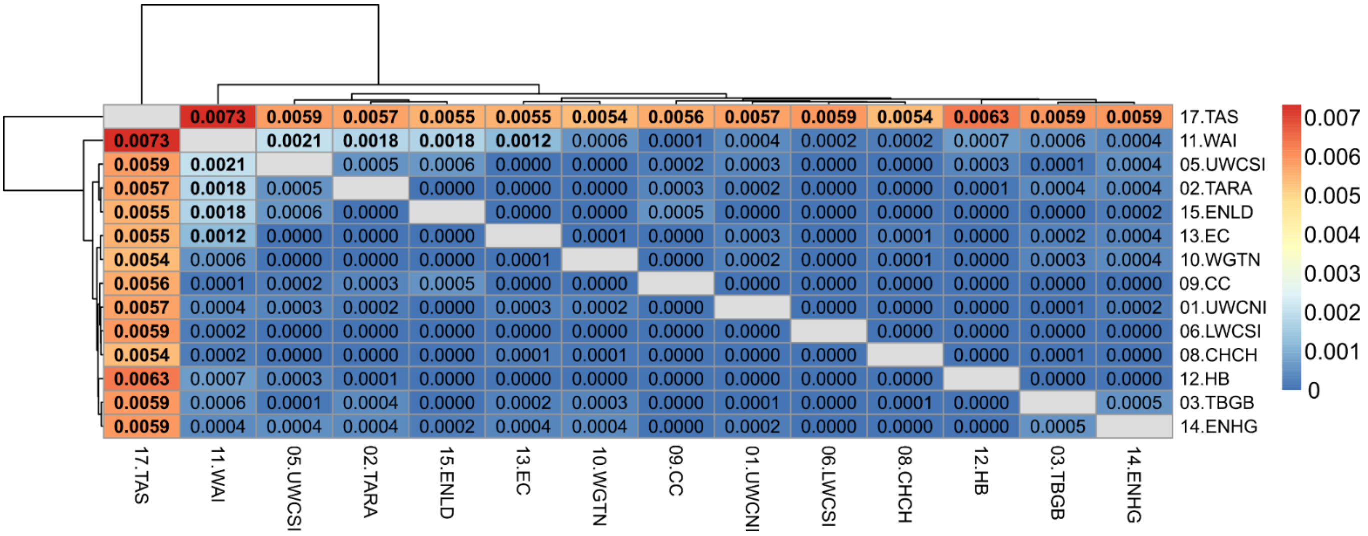


**Supplementary Figure 12.** Heatmap of pairwise weighted $F_{ST}$ estimates (corresponding to the values above and below the diagonal) among sample locations of the neutral SNP dataset. The dendrogram shows the inferred relationship between populations. Significant *P*-values (≤ 0.05 after false discovery rate correction) are in bold.

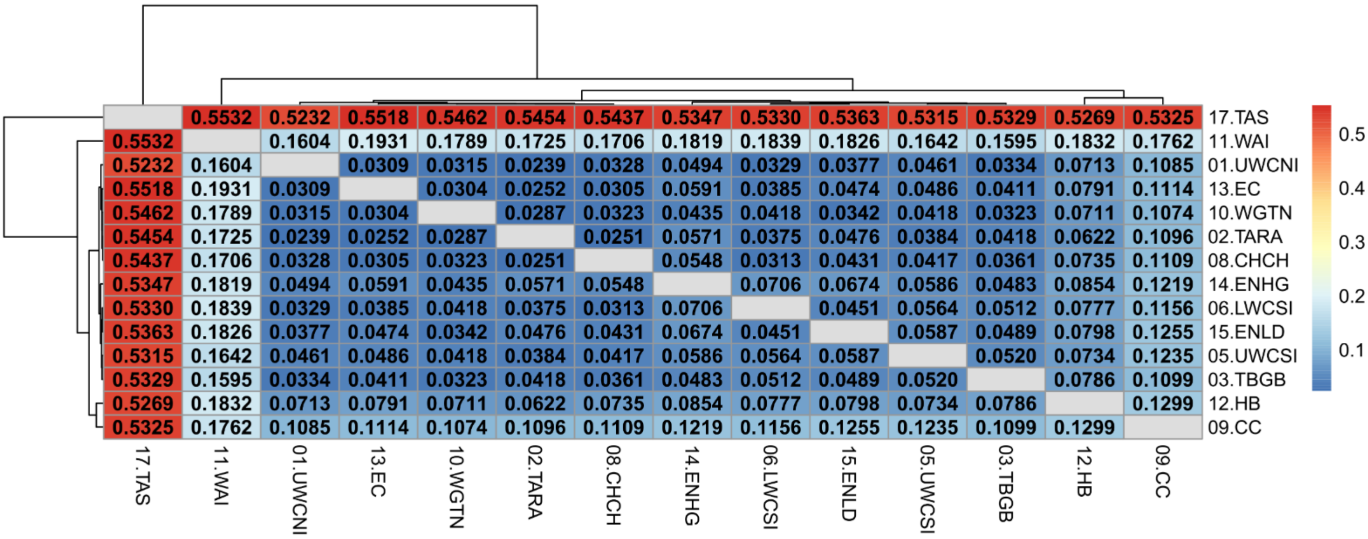


**Supplementary Figure 13.** Heatmap of pairwise weighted $F_{ST}$ estimates (corresponding to the values above and below the diagonal) among sample locations of the adaptive SNP dataset. The dendrogram shows the inferred relationship between populations. Significant *P*-values (≤ 0.01 after false discovery rate correction) are in bold.

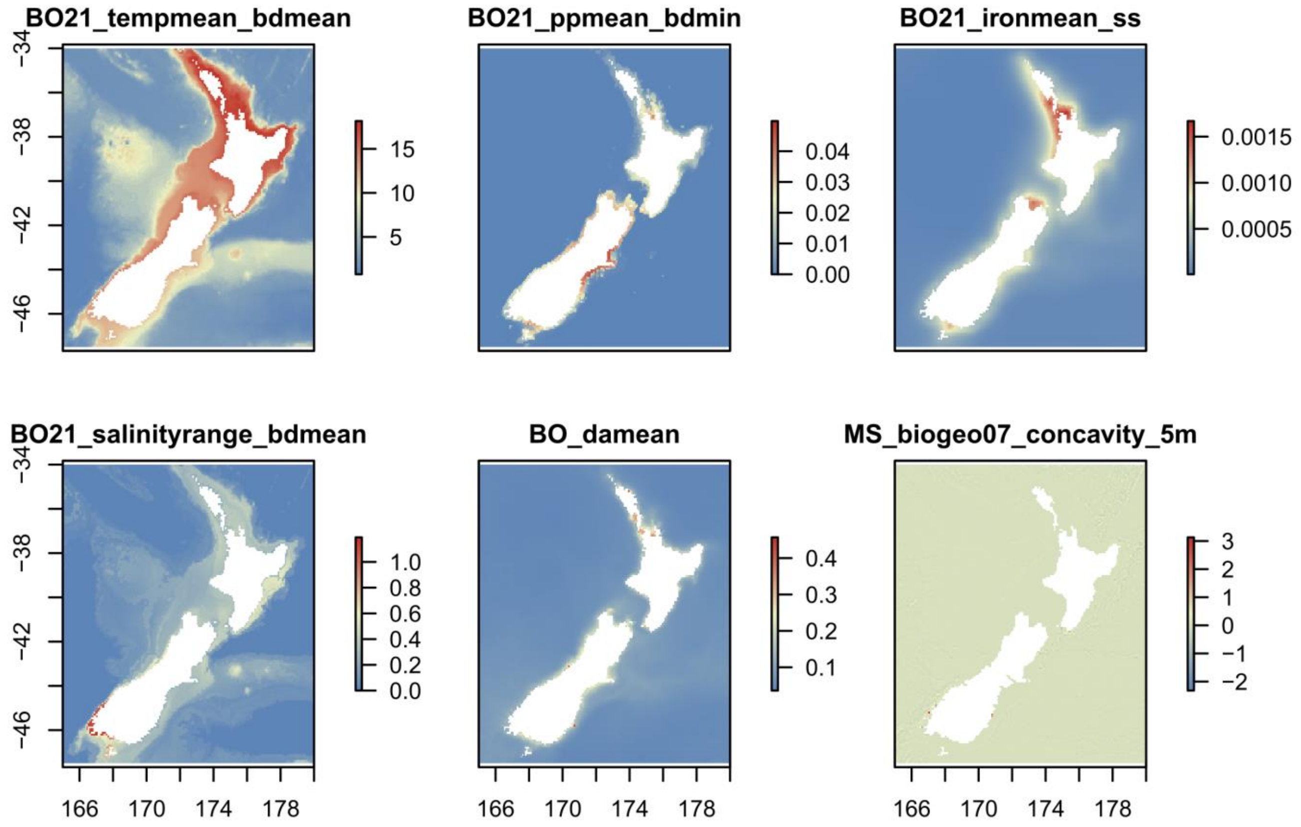


**Supplementary Figure 14.** Plots of values of the six environmental variables used in the Redundancy Analysis (RDA).

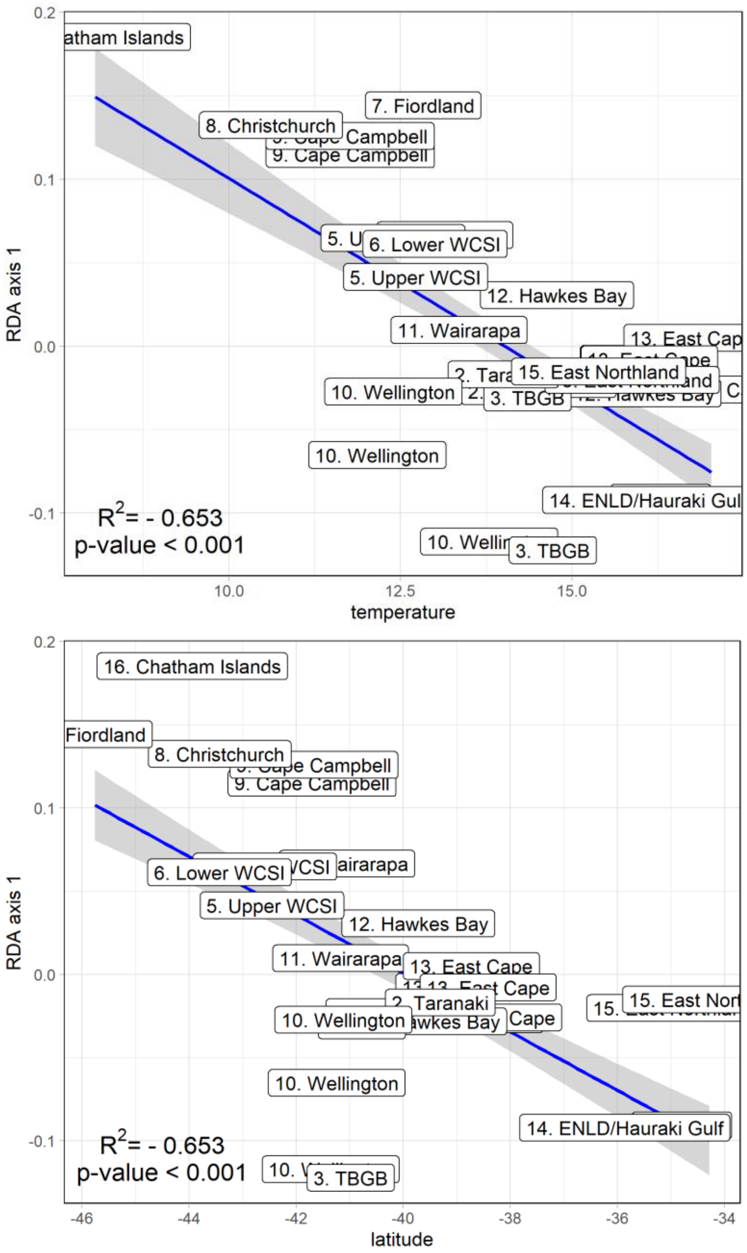


**Supplementary Figure 15.** Linear regression of the axis 1 coordinates of the RDA in relation to mean temperature (top) and latitude (bottom) of sampling locations.

### 1.2 Supplementary Tables

**Supplementary Table 1.** Parameters used in the *K*-mean clustering and DAPC analyses.

| | find.clusters | | | dapc | |
|---|---|---|---|---|---|
| | max.n.clust | n.pca | n.clust | n.pca | n.da |
| Pruned | 100 | 150 | 9 | 50 | 8 |
| Neutral | 100 | 150 | 15 | 50 | 14 |
| Adaptive 1 | 14 | 150 | 6 | 50 | 5 |
| Adaptive 2 | 14 | 150 | 5 | 50 | 4 |

**Supplementary Table 2.** Groups of environmental variables obtained from Bio-ORACLE and MARSPEC.

| Group | *n* variables |
|---|---|
| bathymetry | 6 |
| calcite | 1 |
| carbon phytoplankton biomass | 24 |
| chlorophyll | 22 |
| cloud fraction | 3 |
| current velocity | 24 |
| diffuse attenuation | 3 |
| dissolved oxygen | 25 |
| ice cover | 12 |
| iron concentration | 24 |
| light penetration | 18 |
| nitrate | 25 |
| pH (acidity) | 1 |
| phosphate | 25 |
| photosynthetically available radiation | 2 |
| primary production | 24 |
| salinity | 25 |
| sea surface salinity | 17 |
| sea surface temperature | 20 |
| sea water temperature | 24 |
| silicate | 25 |
| topology (MARSPEC "biogeo") | 7 |

**Supplementary Table 3.** Loci candidates for local adaptation according to the genotype-environment association analysis conducted on scaffold number 1.

| SNP coordinate | BLASTP label / ID | Most correlated variable | Region | Gene name | Annotation (based on BLATSP best match) |
|---|---|---|---|---|---|
| 1051143 | blastp:XP_027142094.1 | mean temperature | gene | LOC104939922 | regulation of nuclear pre-mRNA domain-containing protein 2 isoform X2 [*Larimichthys crocea*] |
| 1507071 | | mean temperature | repeat | | |
| 1586380 | blastp:XP_023122240.1 | primary production | gene | flad1 | FAD synthase [*Amphiprion ocellaris*] |
| 1626325 | blastp:XP_022612259.1 | mean temperature | gene | LOC111230007 | pre-B-cell leukemia transcription factor-interacting protein 1-like [*Seriola dumerili*] |
| 2197375 | blastp:XP_022621898.1 | mean temperature | gene | rad54b | DNA repair and recombination protein RAD54B [*Seriola dumerili*] |
| 2201535 | | mean temperature | none | | |
| 2238075 | blastp:XP_008288143.1 | mean temperature | gene | tmem67 | PREDICTED: meckelin [*Stegastes partitus*] |
| 2364205 | | mean temperature | repeat | | |
| 2798928 | | mean temperature | repeat | | |
| 2879291 | blastp:XP_023148924.1 | mean temperature | gene | mindy3 | ubiquitin carboxyl-terminal hydrolase MINDY-3 [*Amphiprion ocellaris*] |
| 2879525 | blastp:XP_023148924.1 | mean temperature | gene | mindy3 | ubiquitin carboxyl-terminal hydrolase MINDY-3 [*Amphiprion ocellaris*] |
| 3335367 | | mean temperature | none | | |
| 3577629 | blastp:XP_010738915.1 | iron concentration | gene | ankrd28 | serine/threonine-protein phosphatase 6 regulatory ankyrin repeat subunit A isoform X2 [*Larimichthys crocea*] |
| 3991506 | blastp:XP_023138899.1 | mean temperature | gene | LOC111577054 | potassium channel subfamily K member 5-like [*Amphiprion ocellaris*] |
| 4318704 | blastp:XP_008283225.1 | mean temperature | gene | kmt2b | PREDICTED: histone-lysine N-methyltransferase 2B isoform X1 [*Stegastes partitus*] |
| 4504812 | | primary production | repeat | | |
| 4560533 | | mean temperature | none | | |
| 4655135 | blastp:XP_023275606.1 | mean temperature | gene | LOC111664994 | mediator of RNA polymerase II transcription subunit 12-like [*Seriola lalandi dorsalis*] |
| 4655175 | blastp:XP_023275606.1 | mean temperature | gene | LOC111664994 | mediator of RNA polymerase II transcription subunit 12-like [*Seriola lalandi dorsalis*] |
| 6111582 | | salinity range | none | | |
| 6174709 | | mean temperature | none | | |
| 6217015 | | mean temperature | none | | |
| 6404576 | | mean temperature | none | | |
| 6444298 | | mean temperature | repeat | | |

| | | | | | |
|---|---|---|---|---|---|
| 6559859 | | mean temperature | none | | |
| 6640383 | blastp:XP_005450598.1 | mean temperature | gene | e2f3 | transcription factor E2F3 isoform X2 [*Oreochromis niloticus*] |
| 7018261 | TARdn00343 | mean temperature | gene | | Unnanotated gene with multiple isoforms |
| 7132428 | | primary production | none | | |
| 7200389 | blastp:AXY87852.1 | mean temperature | gene | *NA* | glyceraldehyde-3-phosphate dehydrogenase [*Lateolabrax maculatus*] |
| 7333901 | | mean temperature | none | | |
| 7857016 | blastp:XP_008298216.1 | primary production | gene | dedd2 | PREDICTED: DNA-binding death effector domain-containing protein 2 isoform X1 [*Stegastes partitus*] |
| 8485657 | blastp:XP_018527968.1 | iron concentration | gene | lipe | PREDICTED: hormone-sensitive lipase isoform X2 [*Lates calcarifer*] |
| 8559826 | | mean temperature | repeat | | |
| 8720370 | PB.386.8\|TRPT%2F43556 | mean temperature | ? | | Unnanotated single transcript |
| 9111300 | blastp:XP_019122739.1 | mean temperature | gene | ppp1r8 | nuclear inhibitor of protein phosphatase 1 [*Larimichthys crocea*] |
| 9157049 | blastp:XP_018554147.1 | mean temperature | gene | maneal | PREDICTED: glycoprotein endo-alpha-1 2-mannosidase-like protein [*Lates calcarifer*] |
| 9191706 | blastp:XP_019122759.2 | mean temperature | gene | inpp5b | type II inositol 1 4 5-trisphosphate 5-phosphatase isoform X1 [*Larimichthys crocea*] |
| 9603410 | blastp:XP_022620042.1 | mean temperature | gene | oscp1 | protein OSCP1 isoform X3 [*Seriola dumerili*] |
| 9694705 | blastp:XP_010729940.3 | mean temperature | gene | LOC104919580 | thyroid hormone receptor-associated protein 3 isoform X1 [*Larimichthys crocea*] |
| 9763086 | | primary production | none | | |
| 9879614 | | mean temperature | none | | |
| 10423010 | | mean temperature | none | | |
| 10889404 | | mean temperature | none | | |
| 10912349 | blastp:XP_017578417.1 | mean temperature | gene | etfb | PREDICTED: electron transfer flavoprotein subunit beta [*Pygocentrus nattereri*] |
| 11037901 | blastp:XP_010743327.2 | mean temperature | gene | phldb3 | pleckstrin homology-like domain family B member 3 [*Larimichthys crocea*] |
| 11381521 | blastp:XP_010792034.1 | mean temperature | gene | LOC104964847 | PREDICTED: lysosomal thioesterase PPT2-like [*Notothenia coriiceps*] |
| 11455617 | blastp:XP_027142118.1 | mean temperature | gene | LOC104933338 | gamma-aminobutyric acid type B receptor subunit 1 isoform X2 [*Larimichthys crocea*] |
| 11937663 | blastp:XP_022602129.1 | mean temperature | gene | mtmr11 | myotubularin-related protein 11 [*Seriola dumerili*] |
| 12151987 | | mean temperature | repeat | | |
| 12417477 | blastp:XP_008279392.1 | mean temperature | gene | LOC103356862 | PREDICTED: raftlin-like isoform X1 [*Stegastes partitus*] |
| 13020685 | blastp:XP_014876155.1 | mean temperature | gene | LOC106938575 | PREDICTED: septin-7 isoform X2 [*Poecilia latipinna*] |

| | | | | | |
|---|---|---|---|---|---|
| 13170301 | blastp:XP_022056912.1 | mean temperature | gene | thsd7a | thrombospondin type-1 domain-containing protein 7A isoform X3 [*Acanthochromis polyacanthus*] |
| 13299508 | blastp:XP_024241718.1 | mean temperature | gene | LOC112223020 | von Willebrand factor D and EGF domain-containing protein-like [*Oncorhynchus tshawytscha*] |
| 13483672 | | mean temperature | repeat | | |
| 13868517 | blastp:XP_026229042.1 | mean temperature | gene | cspg5a | chondroitin sulfate proteoglycan 5 isoform X1 [*Anabas testudineus*] |

**Supplementary Table 4.** Gene Ontology terms associated to the coding genes from Supplementary Table 3.

| Protein names | Gene names | Gene ontology (GO) |
|---|---|---|
| FAD synthase (EC 2.7.7.2) (FAD pyrophosphorylase) (FMN adenylyltransferase) (Flavin adenine dinucleotide synthase) [Includes: Molybdenum cofactor biosynthesis protein-like region; FAD synthase region] | flad1 zgc:91843 | cytoplasm [GO:0005737]; ATP binding [GO:0005524]; FMN adenylyltransferase activity [GO:0003919]; FAD biosynthetic process [GO:0006747] |
| RAD54 homolog B | rad54b | nucleus [GO:0005634]; ATP binding [GO:0005524]; DNA translocase activity [GO:0015616]; nucleosome-dependent ATPase activity [GO:0070615]; double-strand break repair via homologous recombination [GO:0000724]; reciprocal meiotic recombination [GO:0007131] |
| Transmembrane protein 67 | tmem67 | ciliary transition zone [GO:0035869]; integral component of membrane [GO:0016021]; MKS complex [GO:0036038]; cilium assembly [GO:0060271]; cilium movement [GO:0003341]; convergent extension involved in axis elongation [GO:0060028]; convergent extension involved in gastrulation [GO:0060027]; gastrulation [GO:0007369] |
| Ubiquitin carboxyl-terminal hydrolase MINDY-3 (EC 3.4.19.12) (Deubiquitinating enzyme MINDY-3) (Protein CARP) | mindy3 carp fam188a zgc:153892 | nucleus [GO:0005634]; Lys48-specific deubiquitinase activity [GO:1990380]; thiol-dependent deubiquitinase [GO:0004843]; apoptotic process [GO:0006915] |
| E2F transcription factor 3 | e2f3 | RNA polymerase II transcription regulator complex [GO:0090575]; DNA-binding transcription factor activity, RNA polymerase II-specific [GO:0000981]; protein dimerization activity [GO:0046983]; RNA polymerase II cis-regulatory region sequence-specific DNA binding [GO:0000978]; regulation of transcription by RNA polymerase II [GO:0006357] |
| Protein phosphatase 1, regulatory subunit 8a | ppp1r8a ppp1r8 | nuclear speck [GO:0016607]; mRNA binding [GO:0003729]; protein serine/threonine phosphatase inhibitor activity [GO:0004865]; negative regulation of protein dephosphorylation [GO:0035308] |
| Glycoprotein endo-alpha-1,2-mannosidase-like protein (EC 3.2.1.-) | maneal si:ch211-30b16.2 | Golgi apparatus [GO:0005794]; Golgi membrane [GO:0000139]; integral component of membrane [GO:0016021]; alpha-mannosidase activity [GO:0004559] |

| | | |
|---|---|---|
| Inositol polyphosphate-5-phosphatase B (Fragment) | inpp5b | cytosol [GO:0005829]; membrane [GO:0016020]; inositol-1,4,5-trisphosphate 5-phosphatase activity [GO:0052658]; phosphatidylinositol-4,5-bisphosphate 5-phosphatase activity [GO:0004439]; cilium assembly [GO:0060271]; inositol phosphate dephosphorylation [GO:0046855]; Kupffer's vesicle development [GO:0070121]; melanosome transport [GO:0032402]; phosphatidylinositol dephosphorylation [GO:0046856]; pronephric duct morphogenesis [GO:0039023]; response to epinephrine [GO:0071871] |
| Zgc:171454 protein | oscp1a oscp1 zgc:171454 | cilium assembly [GO:0060271] |
| Electron transfer flavoprotein subunit beta (Beta-ETF) | etfb | mitochondrial matrix [GO:0005759]; electron transfer activity [GO:0009055] |
| Myotubularin-related protein 11 | mtmr11 | phosphatidylinositol-3-phosphatase activity [GO:0004438]; phosphatidylinositol dephosphorylation [GO:0046856] |
| Thrombospondin type-1 domain-containing protein 7A | thsd7aa thsd7a si:dkey-12h3.1 | cell projection [GO:0042995]; integral component of membrane [GO:0016021]; plasma membrane [GO:0005886]; actin cytoskeleton reorganization [GO:0031532]; angiogenesis [GO:0001525]; axon extension [GO:0048675]; blood vessel endothelial cell migration [GO:0043534]; glomerular filtration [GO:0003094]; glomerular visceral epithelial cell development [GO:0072015]; glomerulus development [GO:0032835]; regulation of Notch signaling pathway [GO:0008593]; sprouting angiogenesis [GO:0002040] |
| Chondroitin sulfate proteoglycan 5a | cspg5a | integral component of membrane [GO:0016021]; synapse [GO:0045202]; cell projection morphogenesis [GO:0048858]; nervous system development [GO:0007399]; trans-synaptic signaling, modulating synaptic transmission [GO:0099550] |
